# Adhesion-induced cortical flows pattern E-cadherin-mediated cell contacts

**DOI:** 10.1101/2023.04.11.536411

**Authors:** Feyza Nur Arslan, Édouard Hannezo, Jack Merrin, Martin Loose, Carl-Philipp Heisenberg

## Abstract

Metazoan development relies on the formation and remodeling of cell-cell contacts. Dynamic reorganization of adhesion receptors and the actomyosin cell cortex in space and time play a central role in cell-cell contact formation and maturation. Yet, how this process is mechanistically achieved remains unclear. Here, by building a biomimetic assay composed of progenitor cells adhering to supported lipid bilayers functionalized with E-cadherin ectodomains, we show that cortical Actin flows, driven by the depletion of Myosin-2 at the cell contact center, mediate the dynamic reorganization of adhesion receptors and cell cortex at the contact. E-cadherin-dependent downregulation of the small GTPase RhoA at the forming contact leads to both a depletion of Myosin-2 and a decrease of F-actin at the contact center. This depletion of Myosin-2 causes centrifugal F-actin flows, leading to further accumulation of F-actin at the contact rim and the progressive redistribution of E-cadherin from the contact center to the rim. Eventually, this combination of actomyosin downregulation and flows at the contact determine the characteristic molecular organization, with E-cadherin and F-actin accumulating at the contact rim, where they are needed to mechanically link the contractile cortices of the adhering cells.

## Main

The spatial and temporal regulation of cell-cell adhesion plays a fundamental role in development, growth, and homeostasis^1, 2^. Cadherin adhesion receptors are central components regulating mechanical adhesion between cells and triggering signaling over the cell-cell contact^3–6^. Malfunction of Cadherin-mediated cell-cell adhesion leads to developmental defects, and is a marker of cancerous transformations^7^.

The binding of Cadherins over the contact (*trans*-binding) and clustering of Cadherins at the contact (*cis*-binding) are both thought to drive both cell-cell contact formation and maintenance^8–14^. Moreover, signaling from *trans*-bound Cadherins has been implicated in cell-cell contact expansion by remodeling the actomyosin cell cortex^15–22^. Cadherin-mediated modulation of the actomyosin cortex at the contact has also been associated with an accumulation of Cadherins at the contact rim, where they are needed to mechanically link the contractile cortices of the adhering cells^4, 18, 20, 21, 23^. This rim accumulation of Cadherins has further been shown to be mechanosensitive with tension-induced unfolding of Cadherin adhesion complex components, such as alpha-Catenin and Vinculin, increasing mechanical coupling of the adhesion complex to actomyosin cortex^1, 20, 23–25^. Yet, how the cross-talk of Cadherins and the actomyosin cortex dynamically structure cell-cell contacts remains unknown, mainly due to technical limitations in live imaging of entire contacts with high resolution.

Supported lipid bilayers (SLBs) have been used as an effective assay system for visualizing and analyzing the dynamic molecular rearrangements occurring at specific cell contacts, such as the immunological synapse^26^. Employing such SLB systems for mimicking E-cadherin-mediated cell-cell adhesion showed that E-cadherin mobility constitutes a critical factor for E-cadherin recruitment and Actin architecture at the contact^11, 27^. Still, elucidating how E-cadherin acquires its distinct distribution at the contact requires supplementing these SLB systems with dynamic and high-resolution imaging of contact formation and maintenance.

Here we have combined SLBs as a biomimetic system with high-resolution live imaging to analyze how the dynamic interplay between Cadherins and the actomyosin cortex structures cell contacts. We found that E-cadherin-dependent downregulation of RhoA signaling depletes Myosin-2 and decreases F-actin levels at the forming contact. As a result of this localized downregulation of cortical actomyosin contractility at the contact center, cortical F-actin flows towards the contact rim, taking along E-cadherin and thereby leading to the characteristic molecular organization of the contact, with F-actin and E-cadherin accumulating at the contact rim.

## Results

### Adhesion signaling regulates cortical actomyosin at the contact by modulating RhoA activity

In order to visualize cell contact formation dynamics with high spatiotemporal resolution, we established a biomimetic assay where zebrafish ectoderm progenitor cells adhere to supported lipid bilayers (SLBs), which carry mobile and correctly-oriented zebrafish E-cadherin (Ecad) ectodomains (EcadECD) (Fig.1a and Supp. Fig.1a). Given the importance of Ecad mobility for the proper establishment of Ecad-dependent cell-SLB adhesion^11^, we modulated Ecad mobility in the mainly 1,2-dioleoyl-sn-glycero-3-phosphocholine (DOPC)-composed fluid bilayers. We adjusted the amounts of 1,2-dioleoyl-sn-glycero-3-[(N-(5-amino-1-carboxypentyl)iminodiacetic acid)succinyl] loaded with Ni^2+^ (Ni-NTA-DOGS) and cholesterol, to control Ecad density^28^ and bilayer fluidity^29^, respectively (Supp. Fig.1b). At molar ratios of 4% Ni-NTA-DOGS and 40% cholesterol, we obtained partially-fluid bilayers where tethered EcadECD diffused at 0.34 ± 0.04 µm^2^/s and on which seeded ectoderm progenitors were able to form large (> 15µm diameter) and stable (> 10 min) contacts (Supp. Fig.1c). By contrast, contact formation was strongly reduced on SLBs which lacked EcadECD or when using progenitor cells with reduced endogenous Ecad expression (*cdh1* morphant cells)^21, 30^ (Supp. Fig.1c), further supporting the notion that these bilayers constitute a specific biomimetic assay system to analyze Ecad-mediated cell contact formation.

Using this assay system, we first investigated the initial steps of contact formation by recording time-lapse movies of newly-forming contacts between ectoderm progenitors and EcadECD-decorated SLBs. We used *Tg(cdh1-mlanYFP)* progenitor cells to monitor endogenous Ecad expression^31^, and imaged contact formation using total internal reflection fluorescence (TIRF) microscopy. Consistent with previous findings that Ecad *trans*-dimers accumulate at forming intercellular contacts^11, 32, 33^, we found that the concentration of Ecad at the contact increased within the first 2-3 min post contact initiation, the time which also constitutes the main period of contact expansion (Fig. 1b,c and Supp. Video 1). By contrast, increased Ecad concentrations were not observed in cells seeded on SLBs lacking EcadECD (Supp. Fig. 2a,a’), suggesting that Ecad on both sides of the contact is required for Ecad accumulation at the forming contact.

**Figure 1.**
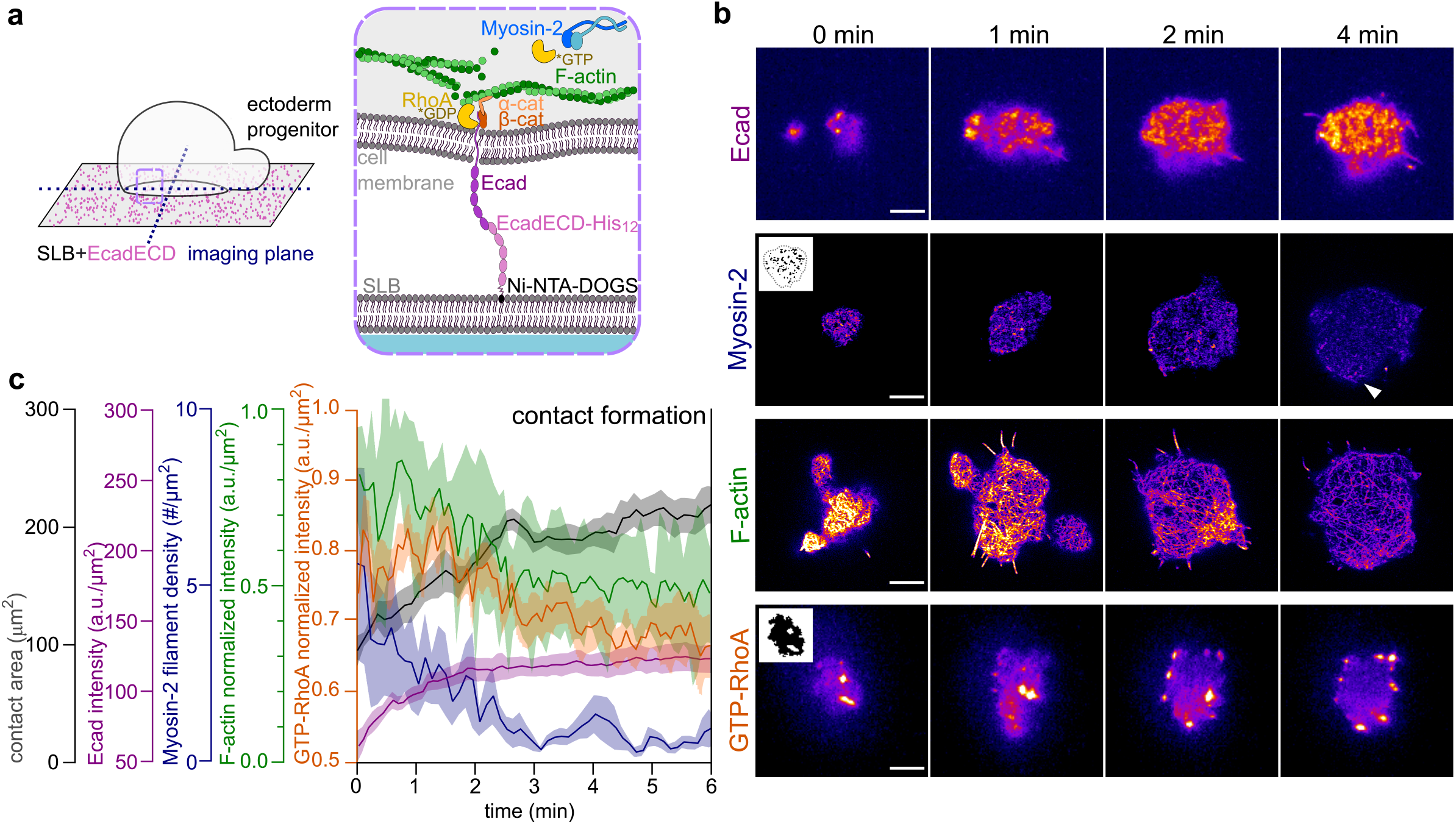
Downregulation of actomyosin and GTP-RhoA coincides with contact area expansion. **(a)** Schematic representations of the biomimetic cell adhesion assay. Left schematic shows the contact interface between the cell and the EcadECD-functionalized SLB, constituting the imaging plane. Right schematic is a close-up of the region marked by the dashed rectangle in the left schematic showing the relevant molecular composition of the contact interface. **(b)** Representative TIRF (for Ecad and GTP-RhoA) or Airyscan (for Myosin-2 and F-actin) contact images of Ecad in a cell obtained from *Tg(cdh1:mlanYFP*), Myosin-2 in a cell obtained from *Tg(actb2:Myl12.1-eGFP)*, F-actin in a cell obtained from Ftractin-mNeonGreen-expressing and GTP-RhoA in a cell obtained from GFP-AHPH-expressing embryos at consecutive steps of contact formation (0, 1, 2 and 4 min post contact initiation). Inlets in Myosin-2 and GTP-RhoA images at 0 min are exemplary masks used for mini-filament density (Myosin-2) and average intensity (GTP-RhoA) calculations shown in (c). White arrowhead at Myosin-2 image at 4 min points to a bleb at the contact interface. Scale bars, 5 µm. **(c)** Plots of contact area (N=4, n=8), Ecad average intensity (N=4, n=10), Myosin-2 mini-filament density (N=4, n=4), F-actin average intensity normalized to maximum intensity (N=6, n=6) and GTP-RhoA average intensity normalized to maximum intensity (N=3, n=11) at the contact as a function of time during contact formation (0-6 min post contact initiation). Data are mean ± s.e.m.

Next, we asked how this accumulation of Ecad at the contact relates to potential changes in the organization of the actomyosin cortex at the contact, to which Ecad couples^3^. Myosin-2 has previously been shown to be reduced at mature homotypic contacts^16, 17, 19, 21^. Using the *Tg(actb2:Myl12.1-eGFP)* progenitor cells^21, 34^ to visualize dynamic changes in Myosin-2 at the forming contact with Airyscan microscopy^35^, we found that cortical Myosin-2, initially decorating the entire contact as mini-filaments, quickly diminished from the contacts during contact area expansion (Fig. 1b,c and Supp. Video 1). This fast reduction in Myosin-2 at the contact led to a nearly complete absence of Myosin-2 at the contact 3 min post contact initiation and depended on the presence of Ecad on both sides of the contact (Supp. Fig. 2a,a’). Despite this general downregulation of Myosin-2 at the contact, some sporadic and short-lived accumulations were still detectable at the contact rim, where cell protrusions such as lamellipodia and blebs retracted (Fig. 1b and Supp. Video 1).

**Figure 2.**
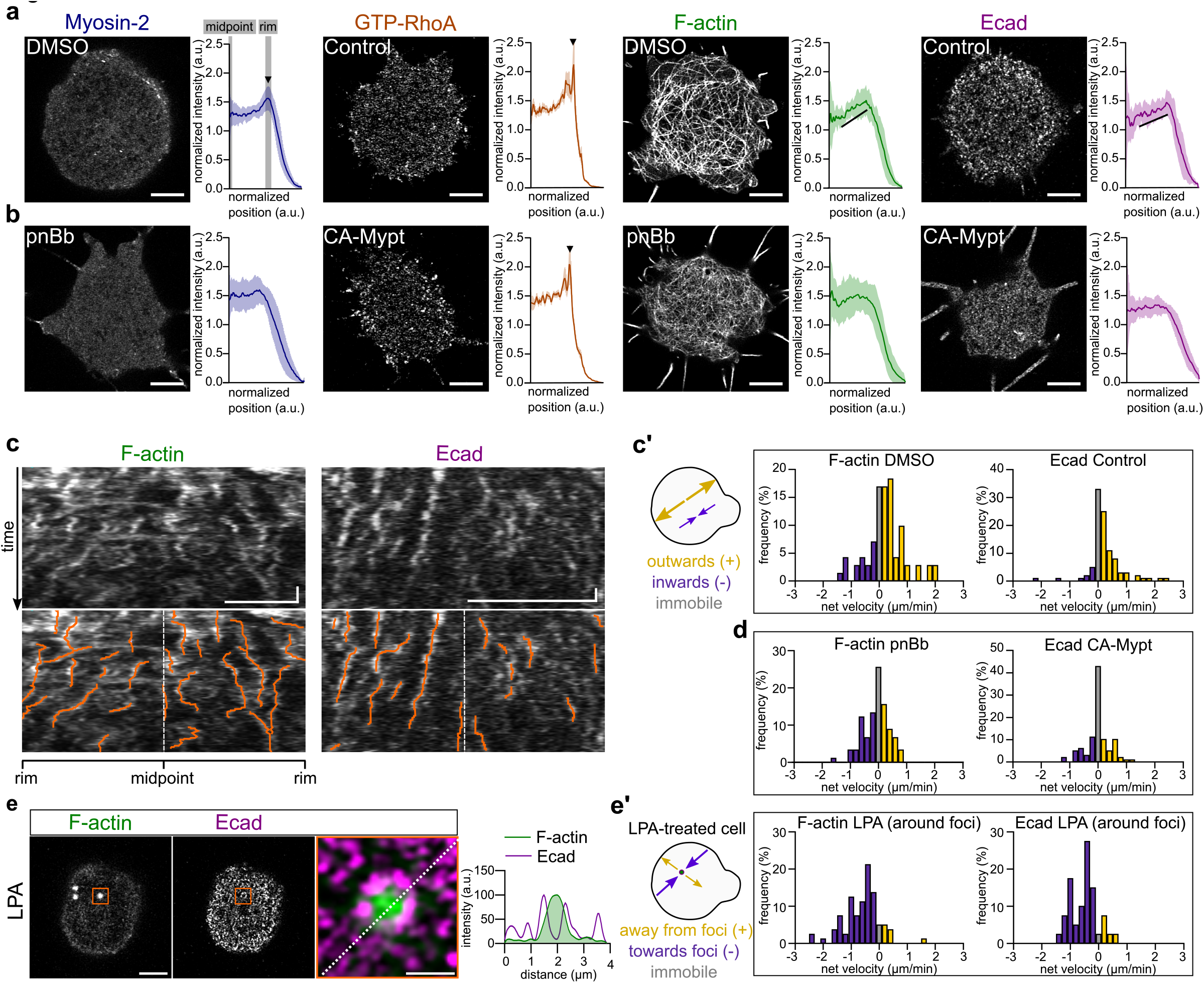
F-actin and Ecad flow towards the contact rim and accumulate there. **(a)** Representative Airyscan images at mature contacts (> 10 min post contact initiation) of Myosin-2 in a cell obtained from *Tg(actb2:Myl12.1-eGFP)*, GTP-RhoA in a cell obtained from GFP-AHPH-expressing, F-actin in a cell obtained from Ftractin-mNeonGreen-expressing and Ecad in a cell obtained from *Tg(cdh1:mlanYFP)* embryos, along with radial intensity plots, normalized to contact length and average intensity, of Myosin-2 (N=3, n=22), GTP-RhoA (N=4, n=20), F-actin (N=3, n=20) and Ecad (N=4, n=20). Black arrowheads in Myosin-2 and GTP-RhoA radial plots point to the intensity peak at the contact rim, black lines in F-actin and Ecad radial plots point to the gradual increase in intensities. For Myosin-2 and F-actin, 0.1% DMSO was added to the medium to control the experimental conditions in (b). Scale bars, 5 µm. Data are mean ± s.e.m. **(b)** Representative Airyscan images at mature contacts of Myosin-2 in a cell obtained from *Tg(actb2:Myl12.1-eGFP)* and F-actin in a cell obtained from Ftractin-mNeonGreen-expressing embryos treated with 10 µM para-nitroBlebbistatin (pnBb), and of Ecad in a cell obtained from *Tg(cdh1:mlanYFP)* and GTP-RhoA in a cell obtained from GFP-AHPH-expressing embryos expressing constitutively active Myosin Phosphatase (CA-Mypt, 70 pg mRNA/embryo), along with radial intensity plots of Myosin-2 (N=3, n=20) and F-actin (N=3, n=24) at contacts of pnBb-treated cells; Ecad (N=3, n=25) and GTP-RhoA (N=3, n=23) in CA-Mypt overexpressing cells. Scale bars, 5 µm. Data are mean ± s.e.m. **(c)** Representative kymographs of F-actin and Ecad flows along the mature contact diameter (top row) in cells obtained from Ftractin-mNeonGreen-expressing or *Tg(cdh1:mlanYFP)* embryos. Detected flow tracks (orange) are superimposed on the raw data (bottom row). Horizontal scale bar, 5 µm; vertical scale bar, 1 min. **(c’)** Histograms of F-actin (N=5, n=7, 70 tracks; mean ± s.d. = 0.22 ± 0.09 µm/min) and Ecad (N=4, n=10, 100 tracks; mean ± s.d. = 0.25 ± 0.17 µm/min) flow velocities at contacts (>3 min post contact initiation), color-coded with yellow for centrifugal/outward-directed tracks, purple for centripetal/inward-directed tracks and gray for immobile tracks (see also schematic on the left). **(d)** Histograms of F-actin and Ecad flow velocities, color-coded as described in (c’), in mature contacts of pnBb-treated (10 µM) (N=7, n=11, 110 tracks; mean ± s.d. = -0.07 ± 0.15 µm/min) or CA-Mypt-expressing cells (70 pg mRNA/embryo) (N=3, n=10, 100 tracks; mean ± s.d. = -0.01 ± 0.13 µm/min) obtained from Ftractin-mNeonGreen-expressing or *Tg(cdh1:mlanYFP)* embryos. **(e)** Representative images of F-actin (left panel) and Ecad (middle panel) at the mature contact of a cell treated with lysophosphatidic acid (LPA, 20 nM) obtained from Ftractin-mKO2-expressing *Tg(cdh1:mlanYFP)* embryos. Higher-magnification dual-color image (right panel) with F-actin in green and Ecad in magenta at a region of the contact (marked by the orange rectangle in left and middle panels), where an ectopic F-actin foci had formed upon LPA treatment. Plot on the right side shows the intensity profiles of F-actin and Ecad along the dashed line shown in the right panel. Scale bars, 5 µm (left and middle panels), 1 µm (right panel). **(e’)** Histograms of F-actin (N=3, n=5, 7 foci, 50 tracks; mean ± s.d. = -0.66 ± 0.79 µm/min) and Ecad (N=3, n=4, 4 foci, 40 tracks; mean ± s.d. = -0.5 ± 0.47 µm/min) flow velocities, color-coded with yellow for tracks directed away from the foci, purple for tracks directed towards the foci and gray for immobile tracks (see also schematic on the left), at regions of the contact in LPA-treated cells where ectopic F-actin foci had formed in cells obtained from Ftractin-mNeongreen-expressing WT or Ftractin-mKO-expressing *Tg(cdh1:mlanYFP)* embryos.

F-actin, similar to Myosin-2, has previously been shown to be partially depleted from the center of mature contacts in several different cell types, including zebrafish ectoderm progenitors^18, 20, 21, 23^. To determine how depletion of Myosin-2 at the contact center during contact expansion is accompanied by dynamic alterations of the Actin cytoskeleton, we analyzed changes in cortical F-actin network organization at the contact of Ftractin-mNeonGreen-expressing cells. This analysis showed that the average F-actin intensity decreased at the contact, when SLBs were decorated with EcadECDs (Fig. 1b,c; Supp. Fig. 2a,a’ and and Supp. Video 1). Notably, this decrease in F-actin intensity at the contact was less pronounced than the observed depletion of Myosin-2 during contact expansion, with some clearly recognizable F-actin cortex still detectable at the mature contact. Collectively, these observations suggest that Ecad contact accumulation is tightly associated with the concomitant reductions in both F-actin and Myosin-2, pointing to the possibility that these processes might be functionally linked.

To determine whether such a functional link exists, we analyzed changes in the activity of the small GTPase RhoA, a critical regulator of both F-actin and Myosin-2^36^, which has previously been suggested to be modulated upon Cadherin binding^18, 19, 37, 38^. To this end, we visualized dynamic changes in RhoA activity during contact formation using an Anillin-based biosensor detecting GTP-RhoA^39, 40^. Similar to Myosin-2 and F-actin, RhoA activity levels decreased at the contact during the first 2-3 min of contact expansion (Fig. 1b,c and Supp. Video 1). By contrast, no such decrease was observed when SLBs were left without EcadECD (Supp. Fig. 2a,a’) or a negative control of the biosensor that does not bind to active RhoA was used^40^ (Supp. Fig. 2b,b’). This concomitant downregulation of RhoA activity with F-actin and Myosin-2 upon Ecad-mediated contact formation suggests that Ecad binding over the contact leads to F-actin and Myosin-2 downregulation at the forming contact by repressing RhoA activity.

To further challenge this suggestion, we analyzed the colocalization of Ecad and GTP-RhoA at the contact. This analysis revealed very little overlap and a negative correlation between Ecad clusters and RhoA activity (Supp. Fig. 2c-c’’), consistent with the notion of Ecad binding leading to local repression of RhoA activity. To further test whether Ecad controls F-actin and Myosin-2 at the contact by modulating RhoA activity, we asked whether the effect of Ecad-mediated contact formation on F-actin and Myosin-2 localization can be overridden by constitutively activating RhoA activity in the contacting cell. To this end, we analyzed contact formation using progenitor cells expressing a constitutively active version of RhoA (CA-RhoA)^41, 42^. We found that while CA-RhoA-expressing progenitors maintained high RhoA activity and increased Ecad at the contact, there was no clearly recognizable reduction of the actomyosin cortex during expansion (Supp. Fig. 2d,d’), supporting the notion that Ecad binding over the contact leads to actomyosin reduction through modulation of RhoA activity.

### Myosin-2 asymmetry leads to centrifugal flows of F-actin and Ecad at the contact

Previous studies have shown that both F-actin and different components of the Cadherin adhesion complex, such as alpha-Catenin, beta-Catenin and Ecad, display a distinct accumulation at the cell contact rim, and that this accumulation is required for mechanically coupling the contractile actomyosin cortices of the adhering cells over the contact^18, 20, 21, 23^. To determine whether such distinct spatial localization can also be observed in our biomimetic assay of contact formation, we first analyzed the average radial distributions of Myosin-2, F-actin, Ecad and GTP-RhoA at mature contacts (> 10 min post contact initiation), a stage where their total intensities remained largely unchanged (Supp. Fig. 2e). Myosin-2 and GTP-RhoA were nearly completely depleted from the contact, except some accumulations at places of the contact rim, where cellular protrusions extended and retracted (Fig. 2a). Averaging this distribution gave rise to Myosin-2 and GTP-RhoA sharply peaking at the contact rim (Fig. 2a). By contrast, F-actin and Ecad displayed a more graded distribution along the contact radius, with increasing levels from the midpoint towards the rim of the contact (Fig. 2a).

In order to dissect the mechanisms by which the actomyosin cortex and adhesion apparatus remodel at the contact, we first determined to what extent this remodeling depends on Ecad binding over the contact. Imaging contacts of cells placed on SLBs lacking EcadECD showed that Ecad, Myosin-2, F-actin and GTP-RhoA at the contact remained homogeneously distributed (Supp. Fig. 3a), suggesting that contact remodeling occurs downstream to Ecad *trans*-binding and signaling. Next, we asked how the specific distributions of Ecad, Myosin-2, F-actin and GTP-RhoA at mature contacts are established during contact formation by monitoring changes in their rim-to-center distributions during Ecad-mediated contact formation. While F-actin, Myosin-2 and Ecad became increasingly localized to the contact rim during the entire contact expansion phase (∼0-3 min post contact initiation), GTP-RhoA levels were sharply downregulated at the contact center already 1 min after contact initiation, with only some active RhoA remaining at the contact rim (Supp. Fig. 3b). This shows that the redistribution and rim accumulation of F-actin, Myosin-2 and Ecad is preceded by a fast downregulation of RhoA activity at the contact center upon contact initiation, pointing to the possibility that suppression of RhoA activity at the contact center leads to the subsequent accumulation of F-actin, Myosin-2 and Ecad at the contact rim.

**Figure 3.**
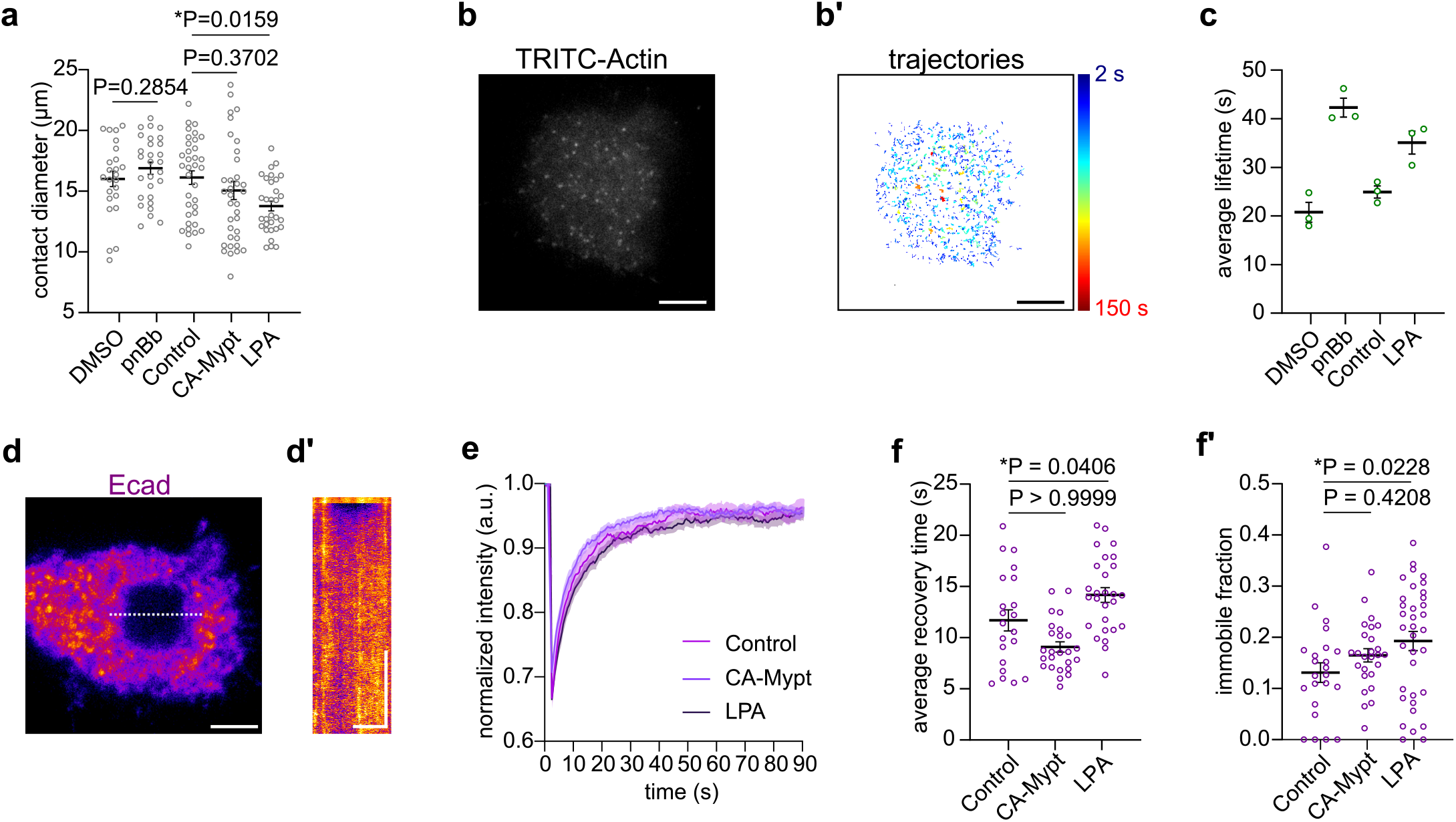
Actomyosin contractility affects contact size and F-actin/Ecad molecular turnover. **(a)** Mature (>10 min post contact initiation) contact diameters of control (DMSO-treated (0.1%) (N=3, n=24) or untreated (N=3, n=35)), para-nitroBlebbistatin-treated (pnBb, 10 μM) (N=2, n=28), constitutively active Myosin Phosphatase-expressing (CA-Mypt, 70 pg mRNA/embryo) (N=3, n=36) and lysophosphatidic acid-treated (LPA, 20 nM) (N=3, n=31) cells. Data are mean ± s.e.m. Student’s t-test for DMSO control and pnBb treatment; Kruskal-Wallis test for untreated control, CA-Mypt expression and LPA treatment. **(b)** Representative TIRF image of TRITC-Actin (0.125 ng/embryo) localization at single-molecule resolution at a mature contact. Scale bar, 5 µm. **(b’)** Time trajectories of detected single TRITC-Actin molecules, as shown in (a), tracked over a period of 6 min. Color map indicates the duration of each trajectory. **(c)** TRITC-Actin average lifetimes at mature contacts for control cells (either treated with 0.1% DMSO (N=3) or untreated (N=3)), and cells treated with either 10 μM pnBb (N=3) or 20 nM LPA (N=3). Each open circle represents a value calculated from 3 different time-lapses obtained on the same day. Data are mean ± s.e.m. **(d)** Representative TIRF image of Ecad at the mature contact of a cell obtained from *Tg(cdh1:mlanYFP)* embryos, at the first post-bleach frame during FRAP measurements. Scale bar, 5 µm. **(d’)** Kymograph of Ecad recovery at the contact, obtained from intensity profiles along the dashed line shown in (d). Horizontal scale bar, 5 µm; vertical scale bar, 1 min. **(e)** Exemplary recovery curves of Ecad intensity within bleached regions normalized to pre-bleach intensity (bleach at 2.5 s) at the mature contacts of cells obtained from *Tg(cdh1:mlanYFP)* embryos for untreated control (N=1, n=11), CA-Mypt-overexpression (70 pg mRNA/embryo) (N=1, n=9) and LPA treatment (20 nM) (N=1, n=9). Data are mean ± s.d. **(f)** Ecad recovery times after photobleaching at mature contacts of untreated control (N=3, n=35), CA-Mypt-expressing (N=3, n=24) and LPA-treated (N=4, n=28) cells obtained from *Tg(cdh1:mlanYFP)* embryos. Data are mean ± s.e.m. Kruskal-Wallis test. **(f’)** Immobile fractions of Ecad at mature contacts, obtained for the data described in (f). Data are mean ± s.e.m. Kruskal-Wallis test.

One possibility by which downregulation of RhoA activity at the contact center might trigger the continuous relocalization of F-actin and Ecad to the contact rim is by building up an actomyosin contractility gradient along the contact radius peaking at the contact rim. This contractility gradient might again give rise to centrifugal F-actin network flows, which, by slowing down at the contact rim, could lead to the rim accumulation of F-actin and Ecad. Consistent with this possibility, we noticed that, although total F-actin intensity at the contact remained unchanged after the initial phase of contact expansion (Fig.1c), F-actin intensity at the contact center continued decreasing for another ∼2 min (up to ∼5 min post contact initiation) (Supp. Fig. 3b,c,c’). This continuous decrease in intensity was accompanied by a decrease in F-actin network density at the contact center and the emergence of an F-actin network density gradient along the contact radius (Supp. Fig. 3c,c’,d), an effect compatible with the possibility of centrifugal F-actin network flows changing F-actin network density along the flow direction^43, 44^. To more directly determine whether the F-actin network indeed displays centrifugal flows at the contact, we performed a kymograph analysis of Actin filament movements at the contact. Strikingly, this analysis revealed persistent centrifugal flows of F-actin at the contact (Fig. 2c,c’; Supp. Fig. 4a,e’ and Supp. Video 2), slowing down at the contact rim (Supp. Fig. 3e). The outward direction of flows was apparent after the initial phase of contact expansion and was also detectable in mature contacts (Supp. Fig. 3f-f’’). Together, this suggests that F-actin network dilution at the contact center and progressive accumulation at the contact rim are achieved and maintained by centrifugal F-actin flows.

To understand how the downregulation of RhoA activity at the contact center might lead to the buildup of an actomyosin contractility gradient and flow along the contact radius, we turned to Myosin-2, previously shown to represent a main determinant of cortical Actin contractility^45–48^. Specifically, we hypothesized that downregulation of RhoA at the contact center might lead to the near complete depletion of Myosin-2 in the center, thereby generating a sharp gradient of Myosin-2 activity along the contact radius (Fig. 2a). To address this hypothesis, we reduced Myosin-2 activity in the contacting cells by exposing cells to the Myosin-2 inhibitor para-nitroblebbistatin (pnBb)^49^ or by expressing a constitutively active form of Myosin Phosphatase (CA-Mypt)^42, 50^ (Fig. 2b; treating cells with pnBb turned out to be not suitable for analyzing GTP-RhoA and Ecad distributions, as the autofluorescence of pnBb strongly decreased their signal-to-noise ratios). Cells exposed to pnBb showed strongly diminished centrifugal F-actin flows (Fig. 2d, Supp. Fig. 4a and Supp. Video 2), and, as a result of this, reduced F-actin network dilution and contact rim accumulation (Fig. 2b and Supp. Fig. 4b). By contrast, pnBb treatment/CA-Mypt expression did not affect the initial signaling-dependent reduction in average F-actin intensity levels during contact formation and concomitant restriction of GTP-RhoA activity to the contact margin (Fig. 2b, Supp. Fig. 4c). This suggests that centrifugal flows of F-actin are predominantly needed for F-actin network density gradient formation and maintenance.

To further explore whether and how the centrifugal flows of F-actin are related to the graded distribution and accumulation of Ecad at the contact rim, we analyzed dynamic changes in Ecad distribution at the contact. Given that Ecad clusters have previously been shown to be taken along by F-actin flows^51–53^, we hypothesized that the observed centrifugal F-actin flows at the contact center might trigger similar flows of Ecad, leading to Ecad gradient formation and contact rim accumulation. Kymograph analysis revealed Ecad clusters to flow centrifugally with an average net velocity similar to F-actin filaments (Fig. 2c,c’, Supp. Fig. 4a and Supp. Video 3), pointing to the possibility that centrifugal F-actin flows take along Ecad towards the contact rim. Consistent with this, distinct Ecad clusters showed partial colocalization with F-actin filaments at the contact (Supp. Fig. 3g-g’’), suggesting that some, but not all Ecad clusters might be directly linked to F-actin. To further challenge the functional link between F-actin and Ecad flows, we tested whether Myosin-2 inhibition not only affects F-actin but also Ecad localization and flows. Blocking Myosin-2 activity with pnBb, while leaving the average intensity of Ecad at the contact unchanged, eliminated Ecad rim accumulation, leading to a homogenous distribution of Ecad across the contact (Supp. Fig. 4c,d), similar to the observations made for F-actin in the presence of pnBb (Fig. 2b and Supp. Fig. 4c). To determine whether the lack of Ecad rim accumulation upon inhibition of Myosin-2 activity is due to reduced centrifugal flows of Ecad, we performed kymograph analysis of Ecad flows in cells with reduced Myosin-2 activity. CA-Mypt-expressing cell contacts not only showed more homogenous Ecad and F-actin intensities and higher F-actin network densities (Fig. 2b and Supp. Fig. 4b,e), similar to the observations made when blocking Myosin-2 activity with pnBb (Fig. 2b and Supp. Fig. 4d) but also displayed strongly reduced centrifugal flows of Ecad clusters (Fig. 2d, Supp. Fig. 4a and Supp. Video 3). This suggests that Ecad clusters might be advected by the F-actin network flows towards the contact rim in a Myosin-2-dependent manner.

**Figure 4.**
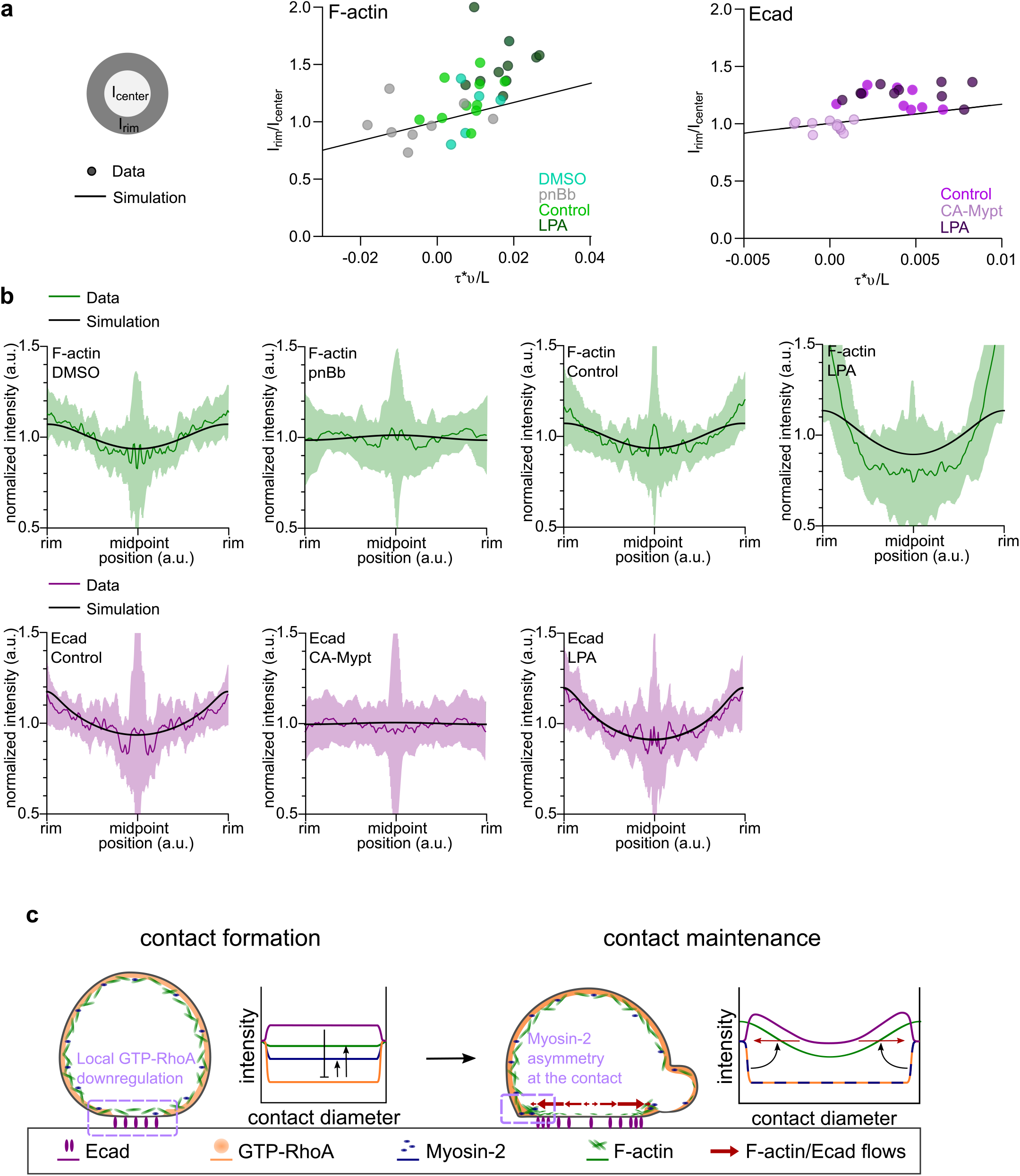
Contractility-dependent flow velocity and molecular turnover determine spatial distribution of F-actin and E-cadherin at the mature contact. **(a)** Plots of rim-to-center intensity ratios for F-actin or Ecad vs parameters predicting rim enrichment (molecular lifetime (τ) * flow velocity (*v*) / contact diameter (L)) across different contractility conditions. Dots represent individual mature (>10 min post initiation) contacts for which rim-to-center intensity ratios (as shown in schematic on the left), flow velocities and contact diameters were measured, and molecular lifetimes were taken from (Supp. Table 1). Lines indicate the predicted values of rim enrichment based on the equation 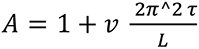 (see Methods). F-actin was imaged in cells obtained from Ftractin-mNeonGreen-expressing embryos (0.1% DMSO (N=4, n=6) as control for 10 µM para-nitroBlebbistatin (pnBb) treatment (N=5, n=8) and untreated control (N=8, n=12) for 20 nM lysophosphatidic acid (LPA) treatment (N=7, n=10)). Ecad was imaged in cells obtained from *Tg(cdh1:mlanYFP)* embryos (untreated control (N=3, n=9) for CA-Mypt-overexpression (70 pg mRNA/embryo) (N=3, n=11) and LPA treatment (N=5, n=10)). (b) Theoretically predicted steady-state F-actin and Ecad intensity profiles based on the equation given in (a) with experimentally measured parameters (Supp. Table 1). For Ecad, an immobile fraction was added to the equation (see Methods). In the same plots, radial F-actin and Ecad intensity plots, normalized to rim-to-rim distance and average intensity, taken from the experiments interfering with cell contractility shown in Fig. 2a,b and Supp. Fig. 4e’,f’ are shown. (c) Schematic of the mechanism defining the molecular patterning of the cell contact. Upon contact initiation *trans*-bound Ecad-dependent adhesion signaling locally downregulates active RhoA, leading to reduction of Myosin-2 and F-actin at the contact center. Consequently, asymmetric distribution of Myosin-2 at the contact drives centrifugal flows of F-actin, along with Ecad, thereby resulting in the accumulation of these proteins at the contact rim.

To further challenge this hypothesis, we asked whether upregulating Myosin-2-dependent actin network contractility is also sufficient for driving F-actin and Ecad centrifugal flows. To this end, we treated cells with lysophosphatidic acid (LPA), which has previously been shown to strongly enhance actomyosin contractility in germ layer progenitor cells^23, 44^. Upon exposure to LPA, F-actin, and to a smaller extent also Ecad, showed an enhanced accumulation at the contact rim, giving rise to steeper gradients of F-actin and Ecad along the radial axis of the contact (Supp. Fig. 4f). Notably, the average intensities of F-actin and Ecad at the contact did not change upon LPA treatment (Supp. Fig. 4c), suggesting that LPA treatment affects the distribution but not the general amount of these proteins at the contact. Unexpectedly, however, radial flow velocities of F-actin and Ecad, and the network density of F-actin at the contact center remained unchanged in LPA-treated contacts compared to untreated controls (Supp. Fig. 4a,b,f’, Supp. Video 2 and Supp. Video 3). Instead, we frequently observed ectopic foci of actomyosin within the contact center (Fig. 2e and Supp. Fig. 4h,i), presumably as a result of high actomyosin network contractility leading to the emergence of local network instabilities and thus the formation of ectopic actomyosin foci, driving local flows of F-actin and Ecad directed towards them^44^ (Fig. 2e’ and Supp. Fig. 4a). Collectively, these findings suggest that polarized distribution of Myosin-2 triggers flows of both F-actin and Ecad, thereby establishing their graded distribution along the radial contact axis and accumulation at the contact rim.

### Cortical flows at the contact determine the contact architecture

To determine whether and how the observed changes in F-actin and Ecad rim accumulations in the different experimental conditions can be explained by centrifugal F-actin flows at the contact, we sought to quantitatively link F-actin flows to F-actin and Ecad rim accumulation. From a theoretical perspective, the strength of this accumulation not only depends on centrifugal flow velocity *v*, but also on contact size *L* and protein turnover/lifetime τ at the contact. A simple conservation equation taking into account these features predicts that at first order, the magnitude of the rim-to-center accumulation *A* should scale as 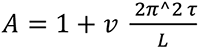. In order to test this quantitatively, we measured contact size and Actin/Ecad lifetimes under different conditions. For contact size, we found that LPA-treated contacts were smaller than untreated controls, consistent with previous observations from germ layer progenitor cell-cell doublets^23^ (Fig. 3a). By contrast pnBb-treated/CA-Mypt-expressing contact sizes, although expected to decrease^23^, remained largely unchanged, likely due to a global decrease in cortical tension leading to cell flattening on the SLB substrate. To measure Actin turnover, we tracked TRITC-Actin protein injected at low amounts using single-molecule imaging at the contact(Fig. 3b,b’). We found a lifetime of ∼25 s for Actin monomers at contacts, which was increased on average by both increasing or decreasing contractility by LPA and pnBb, respectively, in line with previous reports^20, 54^ (Fig. 3c). For Ecad lifetime at the contact, we turned to fluorescence recovery after photobleaching (FRAP) experiments (Fig. 3d,d’,e). Similar to previous observations on alpha-Catenin turnover at Ecad-mediated ectoderm cell-cell contacts^23^, we found an Ecad recovery time of ∼12 s and a small immobile fraction of ∼10%, which were both increased when upregulating actomyosin contractility upon LPA treatment (Fig. 3f,f’). By contrast, decreasing contractility through expression of CA-Mypt had no recognizable effect on Ecad dynamics (Fig. 3f,f’).

Having measured contact size, Ecad/F-actin turnover and flow velocities (Supp. Table 1), we then tested whether the predicted analytical relationship between Ecad/F-actin centrifugal flows and rim accumulation would hold across different contractility conditions and found that the two displayed a robust positive scaling as expected in the analytical expression (Fig. 4a). Explicitly integrating the conservation equation in space (1D along the rim-center axis) and time, starting from a homogenous distribution, showed that contacts reached a steady-state distribution profile in a few min (Supp. Fig. 6a), which closely matched the measured F-actin and Ecad distributions for most experimental conditions (Fig. 4b). Notably, predicted F-actin distributions underestimated the accumulation seen at LPA-treated contacts, suggesting that there might be other factors, in addition to flow velocity, contact size and protein turnover, that also influence F-actin rim accumulations in the presence of LPA. Collectively, this comparison between simulations and experimental observations shows that the progressive accumulation of F-actin and Ecad at the contact rim can, to a large extent, be explained by the centrifugal flows of these proteins at the contact.

## Discussion

Our study identifies a new mechanism by which cell contacts acquire their specific molecular organization required for contact formation and maintenance (Fig. 4c). In particular, we show that centrifugal F-actin flows, triggered by the downregulation of Myosin-2 activity at the contact center, take along Ecad by advection, leading to F-actin and Ecad accumulation at the contact rim, where they are needed for coupling the cortices of the contacting cells. We further show that F-actin and Ecad centrifugal flows depend on the polarized distribution of Myosin-2, which is depleted at the contact center and displays distinct accumulations at places at the contact rim where cellular protrusions retract. Such centrifugal flows could, in principle, be an emergent property of the actomyosin network at the contact, a possibility we tested using an active gel model^55^ (Supp. Fig. 6b). Assuming higher actomyosin contractility at the contact rim as a starting configuration in this model led to long-range Actin network centrifugal flows, similar to those observed experimentally (Supp. Fig. 6b and Methods), which were reinforced and maintained by the resulting accumulation of F-actin at the contact rim. This suggests that the self-organizing properties of the actomyosin network at the contact play an important role in determining the molecular organization of the contact.

Ecad has been previously shown to be transported by F-actin flows, such as at lateral junctions of epithelial cells or during contact initiation between migrating cells^51, 56–58^. F-actin flows, again, are thought to be triggered by the asymmetric distribution of F-actin and/or Myosin-2. Our data suggests that centrifugal F-actin flows at the contact are triggered by the downregulation of RhoA, and as a result of it, Myosin-2 at the contact center. Likely, signaling from *trans*-bound Ecad leads to this downregulation of RhoA activity at the forming contact^37, 38^. Yet, why such Ecad signaling inhibits RhoA and Myosin 2 activity at the contact center but not its rim is unclear. One possibility is that blebs retracting at the contact rim in a Myosin-2-dependent manner^59, 60^ lead to transient accumulations of Myosin-2, thereby establishing the rim-to-center asymmetry of Myosin-2, which is needed for driving centrifugal F-actin flows. Such a mechanism could also explain the observed increased accumulation of actomyosin in LPA-treated cells showing increased blebbing at the contact rim due to their higher contractility^44, 61^ (Supp. Fig. 4f,g). Interestingly, increased accumulation of Myosin-2 at the contact rim of LPA-treated contacts did not lead to faster centrifugal flows (Supp. Fig. 4a, f’). Increased viscosity of the F-actin cortex, for instance, due to decreased turnover and smaller contact sizes^62, 63^, and/or increased friction, for instance, due to more coupling between Ecad and the F-actin cortex^23–25^, might account for F-actin flows not becoming faster despite higher contractility, which we explored in the active gel model (Supp. Fig. 6b and Methods). Additionally, the lower-than-expected velocities might be due to ectopic actomyosin foci forming at the center of contacts (Fig. 2e,e’) and thus interfering with the global centrifugal flow pattern of F-actin. Interestingly, such self-maintained F-actin foci can be recapitulated in active gel models with high contractility^64^.

Another reason why Myosin-2 is specifically downregulated at the contact center and not its rim could be due to Ecad and RhoA/Actin/Myosin-2 accumulation spatially segregating at the rim of mature contacts (Supp. Fig. 3g,h). While RhoA/Actin/Myosin-2 intensities peak at the periphery of the contact, Ecad intensity peaks slightly away from the periphery in a domain directly adjacent to the RhoA/Actin/Myosin-2 accumulations (Supp. Fig. 3h). Such segregation is also apparent at the ectopic foci observed in LPA-treated contacts with Ecad accumulating around the RhoA/Actin/Myosin-2 accumulation in the center of such foci (Fig. 2e and Supp. Fig. 4h,i). Previous studies noted similar spatial segregations of Ecad and F-actin or Ecad and active RhoA^18, 20, 65, 66^, yet the mechanism underlying this segregation remains unclear. It is conceivable that such segregation arises, for instance, by increasing Ecad clustering at the rim altering its binding affinity to the Actin network and thus its transfer range, changes in the actomyosin network architecture making the rim network less susceptible to RhoA/Myosin-2 downregulation by Ecad, and/or membrane height differences between the contact rim and center, due to protrusions or other membrane receptors, spatially restricting Ecad to the contact center^66–70^. Investigating these possible segregation mechanisms will be an interesting subject for future studies.

Mechanosensitivity of Ecad has previously been proposed to also promote Ecad accumulation at the contact rim^20, 23^. Likely, F-actin and Ecad centrifugal flows and Ecad mechanosensation engage in a positive feedback loop, where F-actin centrifugal flows lead to enhanced binding of Ecad to the adjacent Actin network by increasing Actin network density towards the contact rim, allowing the actomyosin network to more efficiently pull on the Ecad adhesion complex. These pulling forces, in turn, might elicit tension-induced conformational changes of the Ecad adhesion complex components alpha-Catenin and/or Vinculin, facilitating coupling of the adhesion complex to the adjacent Actin cortex^24, 25, 71, 72^ and reducing Ecad turnover^20, 23^, thus enhancing Ecad and F-actin accumulation at the rim. Such Ecad and F-actin rim accumulation might then again increase centrifugal F-actin flows, thereby closing a positive feedback loop where flows trigger rim accumulation, and rim accumulation promotes flows^73–75^.

Centrifugal movements of Ecad clusters during contact maturation have previously been noted^20, 32^, but their association with the F-actin cortex remained unclear. Our study, by mechanistically linking the dynamic changes in the F-actin cortex to the redistribution of the Ecad adhesion complex at the contact, provides a generic mechanism by which the maturing contact achieves its specific molecular organization required for contact expansion and maintenance. More generally, it might also explain how cells acquire stable polarity during contact formation by locally remodeling their actomyosin cortex, thereby mechanistically linking cell-cell contact formation to cell polarization in the developing organism.

**Supplementary Figure 1.**
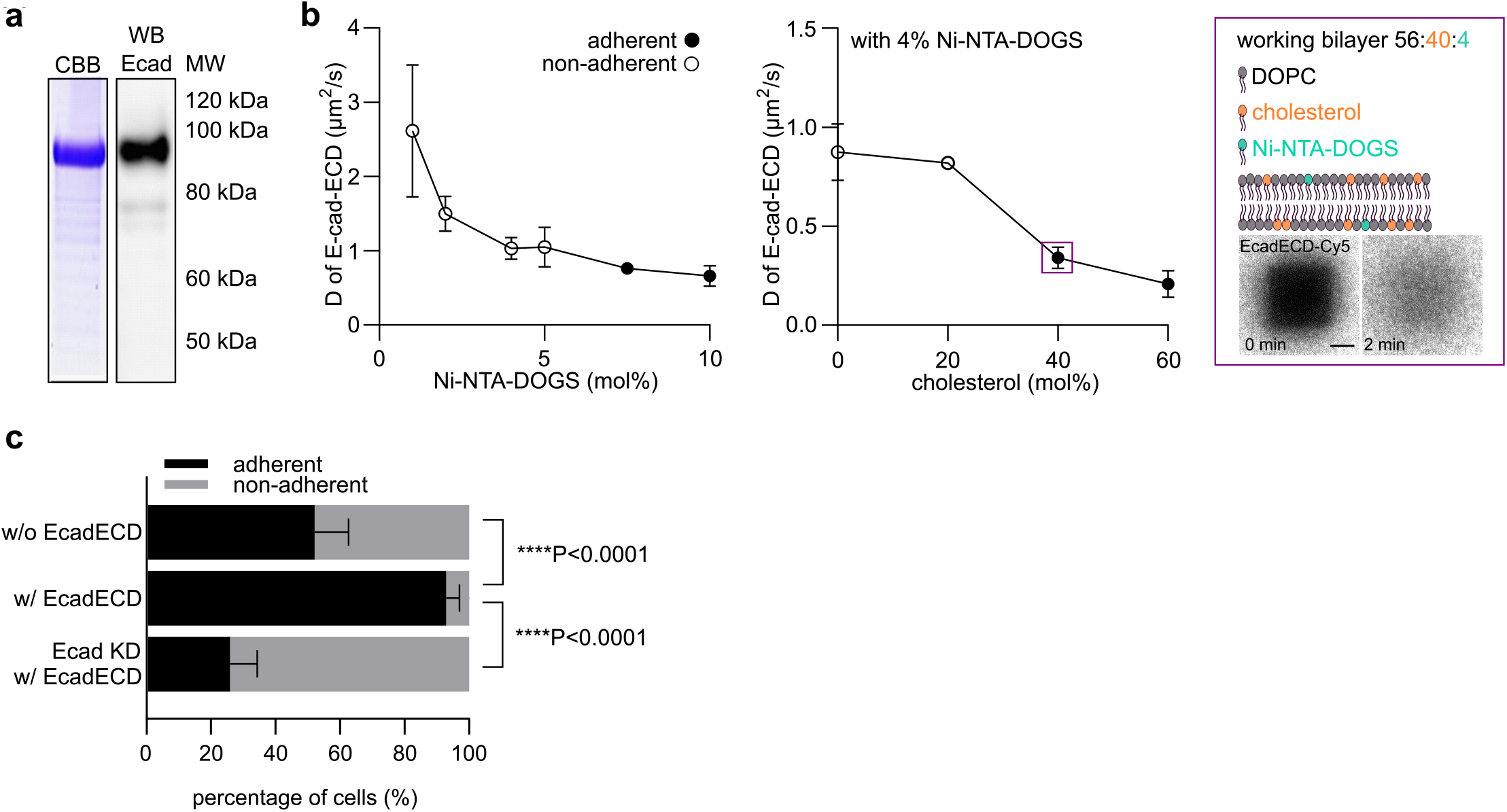
SLB optimization procedures. **(a)** Representative Coomassie Brilliant Blue (CBB) stained SDS-PAGE image (left) and Western blot analysis (right) of eluted zebrafish EcadECD, using anti-zebrafish EcadECD antibody for the western blot. **(b)** Ecad-ECD-Cy5 diffusion constants for different bilayer compositions. Empty circles denote non-adherent (no cell contacts), and full circles adherent (cell contacts) SLBs. Left plot shows the Ecad-ECD-Cy5 diffusion constant as a function of Ni-NTA-DOGS molar concentrations for DOPC-composed SLBs. Middle plot shows Ecad-ECD-Cy5 diffusion constant as a function of cholesterol molar concentrations for DOPC + 4 molar % Ni-NTA-DOGS SLBs. Right schematic shows the bilayer chosen for the experimental work used in the remainder of the study (denoted with a purple rectangle in the middle plot). Below are representative images of a FRAP experiment for Ecad-ECD-Cy5 under this condition (0 and 2 min after bleaching). Scale bar, 5 µm. N=3 independent experiments were performed for each SLB composition. Data are mean ± s.d. **(c)** Percentage of adherent and non-adherent cells on SLBs coated with and without EcadECD (w/o EcadECD N=3, n=48; w/ EcadECD; N=3, n=50) and using Ecad knockdown cells (2 ng *cdh1* morpholino/embryo) (Ecad KD w/ EcadECD; N=3, n=50). Data are mean ± s.e.m. Kruskal-Wallis test.

**Supplementary Figure 2.**
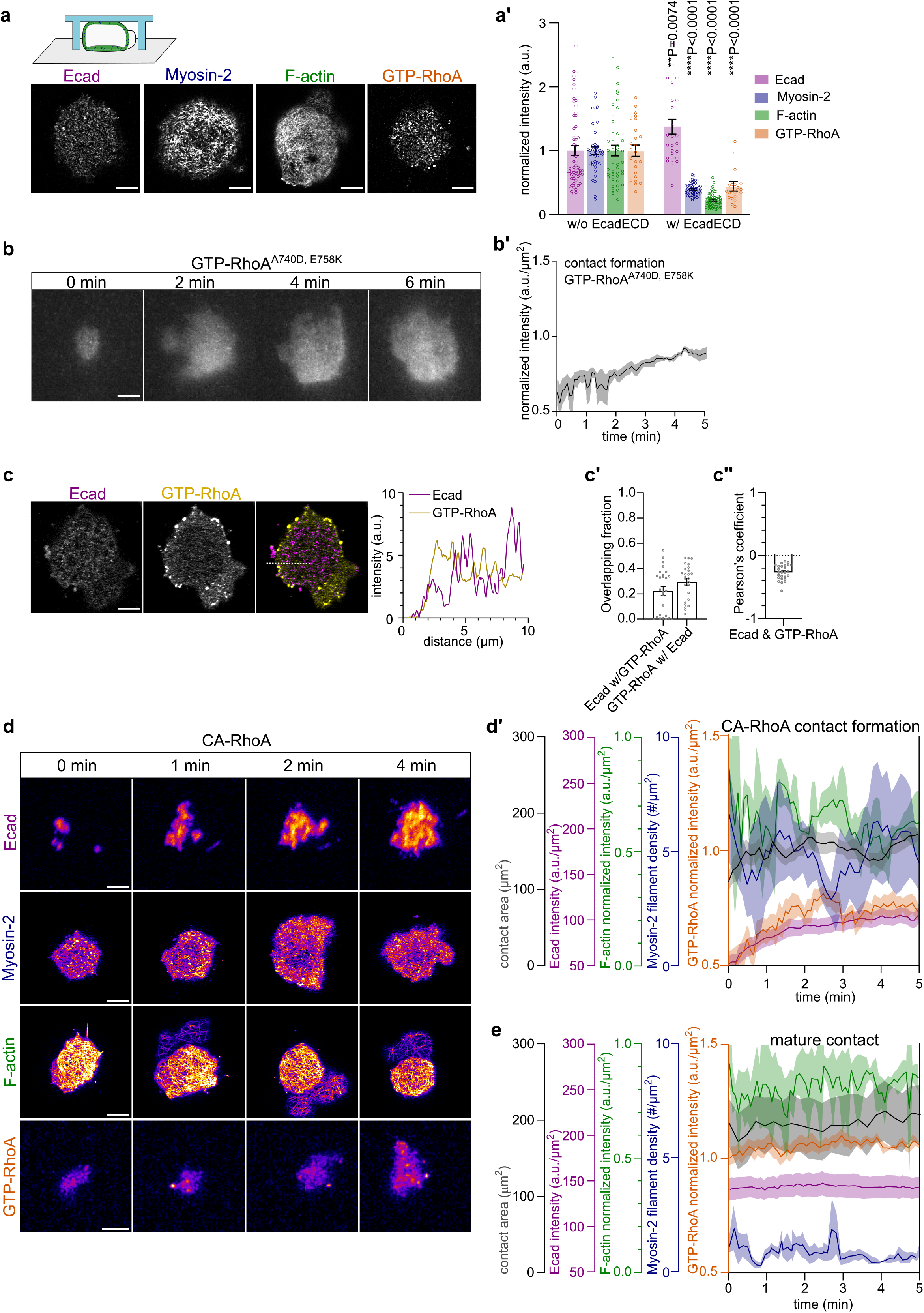
RhoA GTPase and actomyosin activity at cell contacts. **(a)** Representative Airyscan contact images of Ecad in a cell obtained from *Tg(cdh1:mlanYFP)*, Myosin-2 in a cell obtained from *Tg(actb2:Myl12.1-eGFP)*, F-actin in a cell obtained from Ftractin-mNeonGreen-expressing and GTP-RhoA in a cell obtained from GFP-AHPH-expressing embryos, on bilayers lacking EcadECD and spatially confined under a layer of PDMS as shown in the schematic above. Scale bars, 5 µm. **(a’)** Average intensities of Ecad, Myosin-2, F-actin and GTP-RhoA at EcadECD-mediated mature (>10 min post contact initiation) contacts, normalized to average intensities on bilayers lacking EcadECD (imaged as indicated in (a)). Ecad (w/o EcadECD N=3, n=33; w/ EcadECD N=6, n=26), Myosin-2 (w/out EcadECD N=3, n=20; w/ EcadECD N=7, n=50), F-actin (w/out EcadECD N=4, n=23; w/ EcadECD N=3, n=35) and GTP-RhoA (w/out EcadECD N=2, n=25; w/ EcadECD N=4, n=27). Data are mean ± s.e.m. Mann-Whitney *U*-test. **(b)** Representative TIRF images of GTP-RhoA^A740D,E758K^ at contact of a cell obtained fromGFP-AHPH-DM-expressing embryos at consecutive steps of contact formation (0, 2, 4 and 6 min post contact initiation). Scale bar, 5 µm. **(b’)** Plot of GTP-RhoA^A740D,E758K^ average intensity normalized to maximum intensity at the contact as a function of time during contact formation (0-5 min post contact initiation). N=4, n=7. Data are mean ± s.e.m. **(c)** Representative images of Ecad (left panel) and GTP-RhoA (middle panel) at the mature contact of a cell obtained from GFP-AHPH-expressing *Tg(cdh1:tdTomato)* embryos. Dual-color image (right panel) shows Ecad in magenta and GTP-RhoA in yellow. Plot on the right side shows the intensity profiles of Ecad and GTP-RhoA along the dashed line shown in the right panel. Scale bar, 5 µm. **(c’)** Manders’ coefficients M1 (for fraction of Ecad overlapping with GTP-RhoA), M2 (for fraction of GTP-RhoA overlapping with Ecad), and **(c’’)** Pearson’s coefficient (ranging from -1 to 1 indicating the relationship between signal intensities) for colocalization quantification of Ecad and GTP-RhoA. N=2, n=30. Data are mean ± s.e.m. **(d)** Representative TIRF (for Ecad and GTP-RhoA) or Airyscan (for Myosin-2 and F-actin) contact images of Ecad in a cell obtained from *Tg(cdh1:mlanYFP*), Myosin-2 in a cell obtained from *Tg(actb2:Myl12.1-eGFP)*, F-actin in a cell obtained from Ftractin-mNeonGreen-expressing and GTP-RhoA in a cell obtained from GFP-AHPH-expressing embryos expressing constitutively active RhoA (CA-RhoA, 3 pg mRNA/embryo) at consecutive steps of contact formation (0, 1, 2 and 4 min post contact initiation). Scale bars, 5 µm. **(d’)** Plots of contact area (N=3, n=5), Ecad average intensity (N=2, n=13), Myosin-2 mini-filament density (N=3, n=5), F-actin average intensity normalized to maximum intensity (N=3, n=3) and GTP-RhoA average intensity normalized to maximum intensity (N=3, n=9) at the contacts of CA-RhoA-overexpressing cells as a function of time during contact formation (0-5 min post contact initiation). Data are mean ± s.e.m. **(e)** Plots at mature contacts (>10 min post contact initiation) of contact area (N=3, n=4), Ecad average intensity (N=3, n=20) in cells obtained from *Tg(cdh1:mlanYFP)*, Myosin-2 mini-filament density (N=3, n=4) in cells obtained from *Tg(actb2:Myl12.1-eGFP)*, F-actin average intensity normalized to maximum intensity (N=5, n=7) in cells obtained from Ftractin-mNeonGreen-expressing and GTP-RhoA average intensity normalized to maximum intensity (N=4, n=7) in cells obtained from GFP-AHPH-expressing embryos, quantified over 5 min. Data are mean ± s.e.m.

**Supplementary Figure 3.**
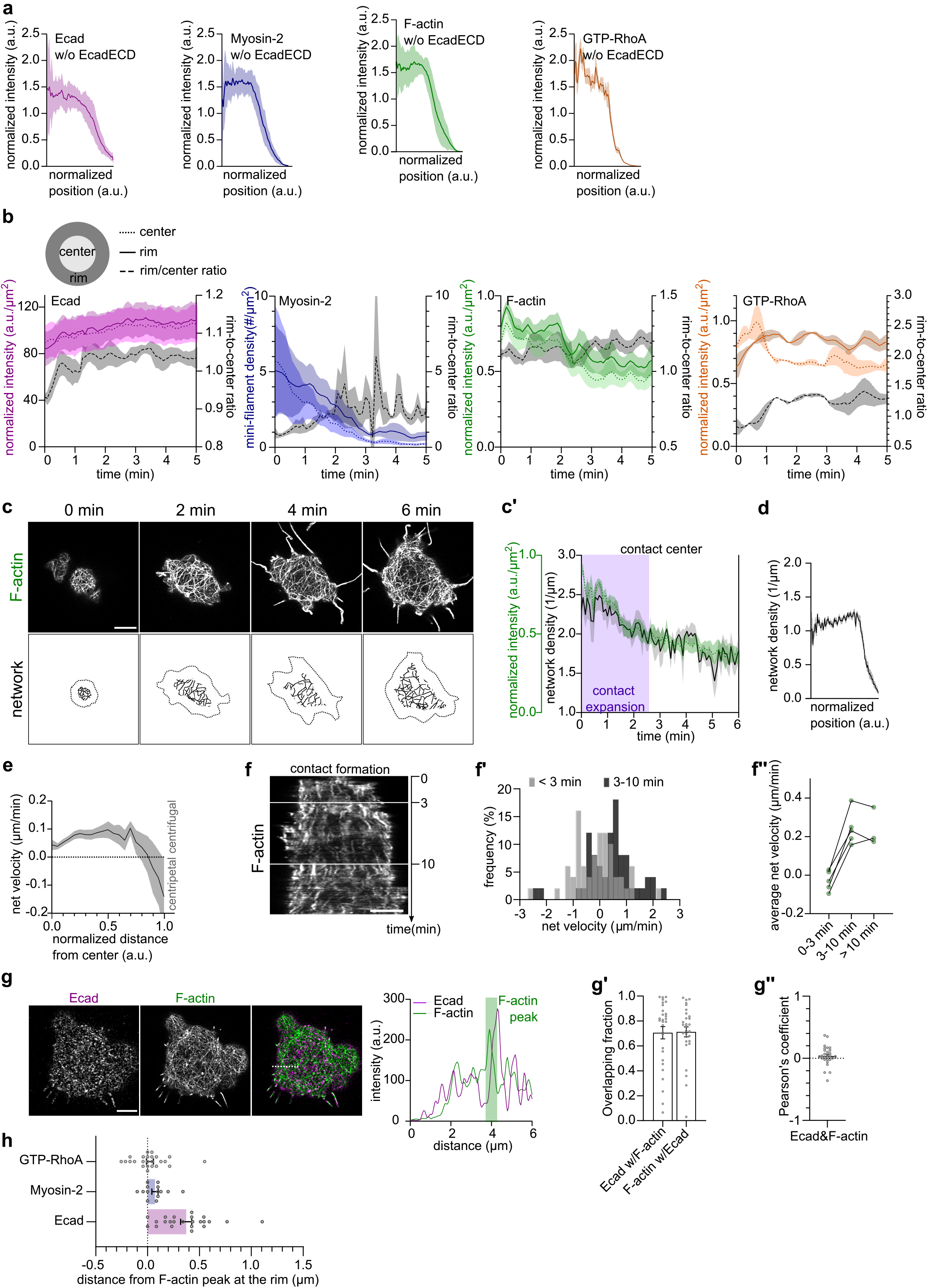
Spatial distribution of adhesion complex/actomyosin proteins and flows during contact formation. **(a)** Radial intensity plots, normalized to contact length and average intensity, at cell contacts on bilayers lacking EcadECD and spatially confined under a layer of PDMS of Ecad (N=3, n=29) in cells obtained from *Tg(cdh1:mlanYFP),* Myosin-2 (N=3, n=18) in cells obtained from *Tg(actb2:Myl12.1-eGFP)*, F-actin (N=3, n=19) in cells obtained from Ftractin-mNeonGreen-expressing and GTP-RhoA (N=2, n=21) in cells obtained from GFP-AHPH-expressing embryos. Data are mean ± s.e.m. **(b)** Plots of average contact rim intensity, average contact center intensity, and rim-to-center intensity ratios of Ecad (N=3, n=5) in cells obtained from *Tg(cdh1:mlanYFP),* Myosin-2 (N=3, n=6) in cells obtained from *Tg(actb2:Myl12.1-eGFP)*, F-actin (N=5, n=5) in cells obtained from Ftractin-mNeonGreen-expressing and GTP-RhoA (N=3, n=3) in cells obtained from GFP-AHPH-expressing embryos as a function of time during contact formation (0-5 min post contact initiation). Data are mean ± s.e.m. **(c)** Representative Airyscan images of F-actin at the contact of a cell obtained from Ftractin-mNeonGreen-expressing embryos during consecutive steps of contact formation (0, 2, 4 and 6 min post contact initiation) (top row) and corresponding images of the skeletonized F-actin networks at contact centers (bottom row). Scale bar, 5 µm. **(c’)** Plots of F-actin average intensity normalized to maximum intensity and corresponding F-actin network density at the contact center as a function of time during contact formation (0-6 min post contact initiation) (N=6, n=6). Purple background marks period of contact expansion. Data are mean ± s.e.m. **(d)** Radial density plot, normalized to contact length, of the F-actin network at mature (>10 min post contact initiation) contacts of cells obtained from Ftractin-mNeonGreen-expressing embryos. N=5, n=23. Data are mean ± s.e.m. **(e)** Radial average flow velocity of F-actin, normalized to contact length, measured along mature contacts of cells obtained from Ftractin-mNeonGreen-expressing embryos. Positive values indicate centrifugal/outward-directed velocities, and negative values indicate centripetal/inward-directed velocities. N=3, n=4. Data are mean ± s.e.m. **(f)** Representative kymograph of F-actin flows during contact formation along the contact diameter of a cell obtained from Ftractin-mNeonGreen-expressing embryos. Scale bar, 5 µm. **(f’)** Histograms of F-actin flow velocities, measured at 0-3 min post contact initiation vs 3-10 min post contact initiation at contacts of cells obtained from Ftractin-mNeonGreen-expressing embryos. N=3, n=5, 50 tracks per time period. **(f’’)** F-actin flow velocities per cell at 0-3, 3-10 and >10 min post contact initiation. **(g)** Representative images of Ecad (left panel) and F-actin (middle panel) at the mature contact of a cell obtained from Ftractin-mKO2-expressing *Tg(cdh1:mlanYFP)* embryos. Dual-color image (right panel) shows Ecad in magenta and F-actin in green. Plot on the right side shows the intensity profiles of Ecad and F-actin along the dashed line shown in the right panel, with green background marking the outermost F-actin peak at the contact rim. Scale bar, 5 µm. **(g’)** Manders’ coefficients M1 (for fraction of Ecad overlapping with F-actin), M2 (for fraction of F-actin overlapping with Ecad) and **(g’’)** Pearson’s coefficient (ranging from -1 to 1 indicating the relationship between signal intensities) for colocalization quantification of Ecad and F-actin. N=2, n=28. Data are mean ± s.e.m. **(h)** Localization of Ecad (N=3, n=24), Myosin-2 (N=3, n=14) and GTP-RhoA (N=2, n=24) intensity peaks in reference to F-actin peaks at contact rims, measured in dual-color contact images of cells obtained from Ftractin-mKO-expressing *Tg(cdh1:mlanYFP), Tg(actb2:Myl12.1-eGFP;actb2:Utrophin-mCherry)* or GFP-AHPH and Ftractin-mKO-expressing WT embryos. Positive values indicate the direction of the contact center, and negative values indicate the direction of the contact rim.

**Supplementary Figure 4.**
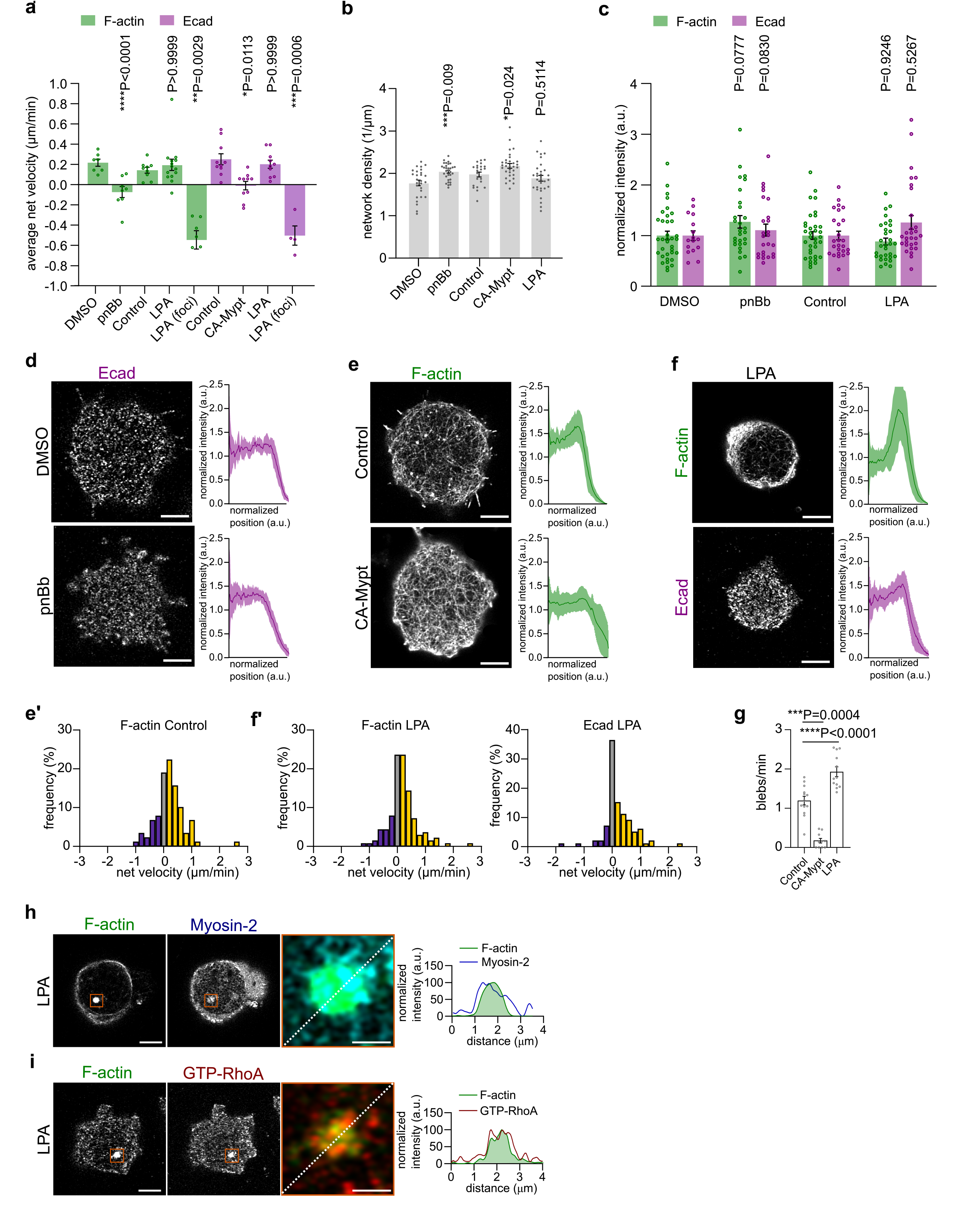
F-actin and Ecad localization and flows at differently contractile cell contacts. **(a)** F-actin and Ecad average flow velocities per cell of F-actin imaged in cells obtained from Ftractin-mNeonGreen-expressing embryos (0.1% DMSO (N=5, n=7) as control for 10 µM paranitroBlebbistatin (pnBb) treatment (N=7, n=11), untreated control (N=6, n=9) for 20 nM lysophosphatidic acid (LPA) treatment (N=7, n=14) and locally around LPA-induced foci (N=3, n=5), and of Ecad imaged in cells obtained from *Tg(cdh1:mlanYFP)* embryos untreated control (N=4, n=10) for 70 pg/embryo constitutively-active Myosin Phosphatase-expressing (CA-Mypt, 70 pg mRNA/embryo) (N=3, n=10), LPA-treated (N=6, n=10) and locally around LPA-induced foci (N=3, n=4). Obtained tracks were also used in construction of histograms displayed in Fig. 2c’,d,e’,(e’) and (f’). Mann-Whitney *U*-test for DMSO control and pnBb. Kruskal-Wallis test for untreated control, LPA and LPA foci. Data are mean ± s.e.m. **(b)** F-actin network density calculated at the mature (>10 min post contact initiation) contact centers of 0.1% DMSO control (N=3, n=27), 10 µM pnBb-treated (N=3, n=25), untreated control (N=3, n=22), CA-Mypt-expressing (70 pg mRNA/embryo) (N=4, n=34) and 20 nM LPA-treated (N=3, n=31) cells obtained from Ftractin-mNeonGreen-expressing embryos. Data are mean ± s.e.m. cells obtained from Ftractin-mNeonGreen-expressing embryos. Data are mean ± s.e.m. Student’s t-test for DMSO control and pnBb. ANOVA test for untreated control, CA-Mypt and LPA. **(c)** Average intensities at mature contacts of F-actin in cells obtained from Ftractin-mNeonGreen-expressing embryos and Ecad in cells obtained from *Tg(cdh1:tdTomato)* (DMSO and pnBb) or *Tg(cdh1:mlanYFP)* (untreated control and LPA) embryos, treated with 0.1% DMSO (F-actin N=3, n=39; Ecad N=2, n=16) as control for 10 µM pnBb treatment (F-actin N=3, n=28; N=2, n=25), or untreated (F-actin N=3, n=35; Ecad N=6, n=26, data is re-used from Supp. Fig. 2a’) as control for 20 nM LPA treatment (F-actin N=3, n=30; Ecad N=4, n=29). Data are mean ± s.e.m. Mann-Whitney *U*-test. **(d)** Representative Airyscan images at mature contacts, of Ecad in cells obtained from *Tg(cdh1:tdTomato)* embryos, along with radial intensity plots, normalized to contact length and average intensity, for 0.1%DMSO (N=2, n=15) and 10 µM pnBb (N=2, n=24) treatments. Scale bars, 5 µm. Data are mean ± s.e.m. **(e)** Representative Airyscan images at mature contacts, of F-actin in cells obtained from Ftractin-mNeonGreen-expressing embryos, along with radial intensity plots, normalized to contact length and average intensity, for untreated control (N=3, n=31) and CA-Mypt-expressing cells (70 pg mRNA/embryo) (N=2, n=25). Scale bars, 5 µm. **(e’)** Histogram of F-actin (N=6, n=9, 90 tracks) flow velocities at mature contacts, color-coded with yellow for centrifugal/outward-directed tracks, purple for centripetal/inward-directed tracks and gray for immobile tracks (see also schematic in Fig. 2e’). Data are mean ± s.e.m. **(f)** Representative Airyscan images at 20 nM LPA-treated mature contacts, of F-actin in cells obtained from Ftractin-mNeonGreen-expressing and of Ecad cells obtained from *Tg(cdh1:mlanYFP)* embryos, along with radial intensity plots, normalized to contact length and average intensity. F-actin N=3, n=34 and Ecad N=3, n=16. Scale bars, 5 µm. Data are mean ± s.e.m. **(f’)** Histograms of F-actin (N=7, n=14, 140 tracks) and Ecad (N=6, n=10, 100 tracks) flow velocities at 20 nM LPA-treated mature contacts, color-coded as described in (e’). **(g)** Frequency of blebs in untreated control (N=3, n=13), CA-Mypt-expressing (75 pg/embryo) (N=3, n=10) and 20 nm LPA-treated (N=4, n=12) mature contacts. Data are mean ± s.e.m. ANOVA test. **(h)** Representative images of F-actin (left panel) and Myosin-2 (middle panel) at the mature contacts of a cell treated with 20 nM LPA, obtained from *Tg(actb2:Myl12.1-eGFP;actb2:Utrophin-mCherry)* embryos. Higher-magnification dual-color image (right panel) with F-actin in green and Myosin-2 in dark blue at a region of the contact (marked by the orange rectangle in left and middle panels), where an ectopic F-actin foci had formed upon LPA treatment. Plot on the right side shows the intensity profiles of F-actin and Myosin-2 along the dashed line shown in the right panel. Scale bars, 5 µm (left and middle panels), 1 µm (right panel). **(i)** Representative images of F-actin (left panel) and GTP-RhoA (middle panel) at the mature contacts of a cell treated with 20 nM LPA, obtained from GFP-AHPH and Ftractin-mKO-expressing embryos. Higher-magnification dual-color image (right panel) with F-actin in green and GTP-RhoA in red at a region of the contact (marked by the orange rectangle in left and middle panels), where an ectopic F-actin foci had formed upon LPA treatment. Plot on the right side shows the intensity profiles of F-actin and GTP-RhoA along the dashed line shown in the right panel. Scale bars, 5 µm (left and middle panels), 1 µm (right panel).

**Supplementary Figure 5.**
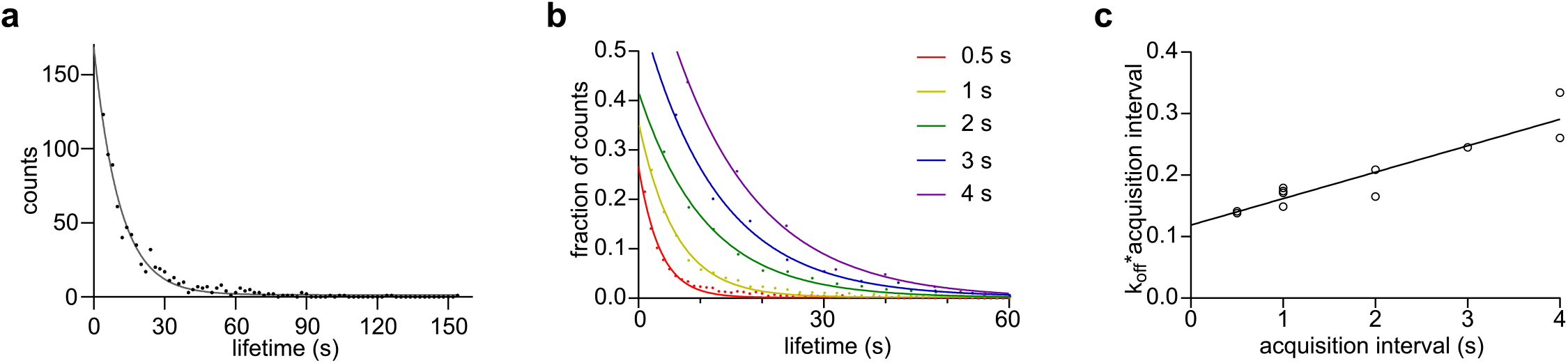
Single-molecule measurement of Actin lifetime. **(a)** Exemplary decay curve of TRITC-Actin single-molecule lifetime distributions from a single cell contact (dots), imaged with 1 s acquisition intervals, overlaid with the best-fit monoexponential decay function (line). n=795 tracks. **(b)** Decay curves of TRITC-Actin single-molecule lifetime at cell contacts imaged with different acquisition intervals (0.5, 1, 2, 3 and 4 s). Dots represent the measurements and lines indicate the monoexponential best fits. (0.5 s N=1, n=2; 1 s N=2, n=4; 2 s N=2, n=2; 3 s N=1, n=1; 4 s N=1, n=2). **(c)** Plot of effective dissociation rates multiplied with acquisition intervals as a function of acquisition interval, corresponding to the data shown in (b). Open circles represent the measurements and the line indicates the best linear fit used for calculating the bleach-corrected dissociation rate.

**Supplementary Figure 6.**
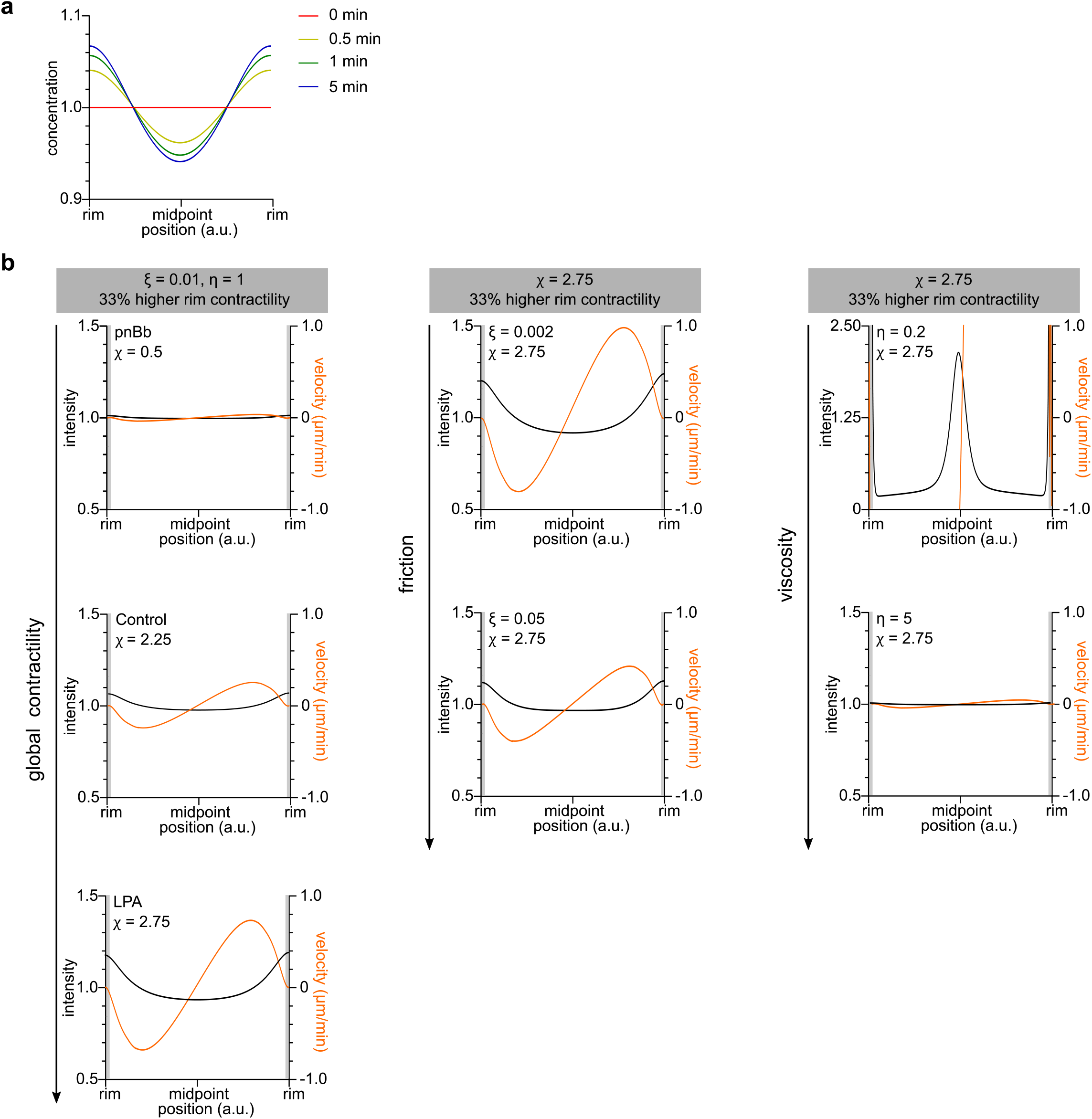
Modeling flows at the contact. **(a)** Theoretically predicted intensity profiles at 0, 0.5, 1 and 5 min for an exemplary condition with protein lifetime of 0.5 min and average flow velocity of 0.2 µm/min at a contact of 20 µm diameter, based on first-order kinetics and the equation given in Fig. 4a. **(b)** Theoretically predicted steady-state intensity and velocity profiles along contacts, based on active gel equations (see Methods). For an increased value of local contractility at the contact rim, increased values of global actomyosin contractility of the gel *χ* (left panel) represent differing contractility conditions in experimental data. Increased values of friction ξ (middle panel) and viscosity η (right panel) are explored for the high global contractility condition.

**Supplementary Table 1.** Experimentally obtained simulation parameters. Average contact diameters (Fig. 3a), flow velocities (Fig.2 c’,d,e’, Supp. Fig. 4e’,f’) and molecular turnover times (Fig. 3c,f,f’) for differing contractility conditions.

|  | Contact diameter (μm) | Actin average flow velocity (μm/min) | Actin average lifetime (s) | Ecad average flow velocity (μm/min) | Ecad average recovery time (s) | Ecad immobile fraction |
| --- | --- | --- | --- | --- | --- | --- |
| **DMSO control** | 16.01 ± 3.11 | 0.25 ± 0.69 | 20.77 ± 2.92 |  |  |  |
| **pnBb** | 16.89 ± 2.64 | -0.07 ± 0.44 | 42.3 ± 2.76 |  |  |  |
| **Untreated control** | 16.12 ± 3.28 | 0.21 ± 0.49 | 24.9 ± 1.8 | 0.25 ± 0.17 | 11.69 ± 4.71 | 0.13 ± 0.09 |
| **CA-Mypt** | 15.06 ± 4.49 |  |  | -0.01 ± 0.13 | 9.10 ± 2.51 | 0.16 ± 0.07 |
| **LPA** | 13.77 ± 2.24 | 0.22 ± 0.57 | 35.1 ± 3.3 | 0.20 ± 0.13 | 14.17 ± 3.73 | 0.19 ± 0.11 |

## Video Legends

**Supplementary Video 1. Contact formation.**

Part 1: Time-lapse TIRF movie of Ecad at the forming contact (0-6 min post contact initiation) of a cell obtained from a *Tg(cdh1:mlanYFP)* embryo. Part 2: Time-lapse Airyscan movie of Myosin-2 at the forming contact of a cell obtained from a *Tg(actb2:myl12.1-eGFP)* transgenic embryo. Part 3: Time-lapse Airyscan movie of F-actin at the forming contact of a cell obtained from an embryo expressing Ftractin-mNeonGreen. Part 4: Time-lapse TIRF movie of GTP-RhoA at the forming contact of a cell obtained from an embryo expressing GFP-AHPH. Left panel: raw movie. Right panel: the raw movie masked to remove bright GFP-AHPH foci. Scale bars, 5 µm.

**Supplementary Video 2. F-actin centrifugal flows at the contact.**

Time-lapse Airyscan movies of F-actin at the mature contacts (>10 min after contact initiation) for the different contractility conditions, with cells obtained from embryos expressing Ftractin-mNeonGreen. Part 1: Contact of an untreated control cell. Part 2: Contact of a cell treated with 10 µM para-nitroBlebbistatin (pnBb). Part 3: Contact of a cell treated with 20 nM lysophosphatidic acid (LPA). The yellow rectangles demarcate ROIs shown with temporal-color coding at the end of the movie. Scale bars, 5 µm for movies and 1 µm for ROIs.

**Supplementary Video 3. E-cadherin centrifugal flows at the contact.**

Time-lapse Airyscan movie of Ecad at the mature contacts (>10 min after contact initiation) for the different contractility conditions, with cells obtained from *Tg(cdh1:mlanYFP)* embryos. Purple circles mark Ecad clusters in randomly selected trajectories of minimum 1 min. Part 1: Contact of an untreated control cell. Part 2: Contact of a cell obtained from an embryo expressing constitutively active Myosin phosphatase (CA-Mypt, 70 pg mRNA/embryo). Part 3: Contact of a cell treated with 20 nM lysophosphatidic acid (LPA). Scale bars, 5 µm.

## Materials and Methods

### Expression and purification of EcadECD

cDNA encoding the zebrafish E-cadherin ectodomain (Q90Z37_DANRE, EC1 to EC5, residues G141 to D672), with an N-terminal human CD33 signaling peptide and C-terminal 12xHis purification tag was codon optimized and ordered as a gBlocks Gene Fragment (IDT) with overhangs for Gibson assembly (NEB). The sequence was engineered to have a single Cys in the EC5 domain for site-specific labeling as previously described^11^. The product was inserted between EcoRI and XbaI sites of pcDNA3.1(-) mammalian expression vector (Invitrogen). EcadECD was expressed in suspension FreeStyle 293-F cells (Thermo Fisher Scientific) in Freestyle 293 Expression Medium at 37°C with 8% CO_2_. FreeStyle 293-F cells were transiently transfected using polyethyleneimine (Polysciences, #23966) in Opti-MEM Reduced Serum Medium (Thermo Fisher Scientific). Cultures were supplemented with 5 mM CaCl_2_ 2 days after transfection^76^, and culture media containing secreted EcadECD was collected 5 days later. Filtered and conditioned media was loaded to HisTrap Fast Flow Crude Column (Cytiva) for affinity chromatography on an ÄKTA Pure Fast Protein Liquid Chromatography system (Cytiva) and eluted with an imidazole gradient at 4°C. Clean fractions, determined by SDS-PAGE stained with Coomassie Brilliant Blue, were pooled together and dialyzed overnight in storage solution (100 mM NaCl, 20 mM Tris-Cl pH 8, 3 mM CaCl_2_). Alternatively, the buffer was exchanged using PD-10 desalting columns (Cytiva). The identity of the protein was verified with N-terminal sequencing. Clean protein was aliquoted at a final 50 µg/ml concentration and snap-frozen for long-term storage at -80°C with 5% glycerol.

### Fluorescent labeling of EcadECD

To perform FRAP experiments for determining the protein diffusion constant, EcadECD was labeled at the Cys residue using Sulfo-C5-maleimide (Lumiprobe). First, the sample was incubated for 20 min with TCEP (100 molar fold excess of protein) at room temperature. Then, maleimide dye (10 molar fold excess of protein) was added to the sample and incubated at room temperature for 1 h. Excess dye was removed using a 7K MWCO Zeba™ Spin Desalting Column (Thermo Fisher Scientific).

### Western blot

Eluted protein fraction was incubated at 70°C in NuPAGE LDS Sample Buffer and NuPAGE Sample Reducing Agent (Invitrogen) for 10 min before getting loaded to a 4-12% NuPAGE Bis-Tris protein gel. After SDS-PAGE, proteins were transferred to a membrane using the iBlot Western Blotting System (Invitrogen) according to the manufacturer’s protocol. For the immunodetection of EcadECD, membrane was blocked with blocking buffer (3% BSA, 0.2% Tween-20 in PBS) for 1 h at room temperature and incubated overnight with rabbit anti-zebrafish Ecad antibody^21^ (1:5000) in blocking solution. After 3x10 min washes with PBT (PBS with 0.2% Tween), membrane was incubated with Peroxidase AffiniPure Goat Anti-Rabbit IgG (H+L) (1:20000) (Jackson ImmunoResearch Laboratories, Inc.) for 45 min at room temperature and washed 4x5 min with PBT, then 2x5 min with PBS. The membrane was developed with Clarity Western ECL Substrate (Bio-Rad) before imaging.

### Preparation of SLBs and functionalization with EcadECD

To make small unilamellar vesicles, 1,2-dioleoyl-sn-glycero-3-phosphocholine (DOPC) (Avanti Polar Lipids), 1,2-dioleoyl-sn-glycero-3-[(N-(5-amino-1-carboxypentyl)iminodiacetic acid)succinyl] (nickel salt, Ni-NTA-DOGS) (Avanti Polar Lipids), 1,2-distearoyl-sn-glycero-3-phosphoethanolamine-N-[amino(polyethylene glycol)-2000] (DSPE-PEG2000) (Avanti Polar Lipids), and freshly dissolved cholesterol (Sigma Aldrich) lipid mixtures in chloroform with a molar ratio of 55.9:4:0.1:40 (unless otherwise stated) were prepared in glass vials and evaporated under N_2_ stream to get a homogenous thin film. Lipids were further vacuumed for 1 h to remove the remaining solvent and frozen at -20°C unless freshly used. Lipid films were resuspended in vesicle buffer (75 mM NaCl, 20 mM Hepes) at 37°C by vortexing to a final concentration of 1.5mM and freeze-thawed in liquid nitrogen 5x before aliquoting. Aliquots were kept at -20°C and used within 2 weeks. For experiments, solutions were diluted to 0.2 mM with vesicle buffer and bath sonicated for 15 min.

To form the lipid bilayers on coverslips, 24×50 mm high-precision coverslips (no. 1.5H; Marienfeld) were cleaned in Piranha solution (3:1, 98% H_2_SO_4_ (Merck):30% H_2_O_2_ (Sigma-Aldrich)) for 1 h. The coverslips were further washed with deionized water and kept in water to be used within 2 weeks. Before use, coverslips were dried, PCR tubes, with their conical ends removed, were attached to the coverslips as reaction chambers, using ultraviolet curing glue (Norland optical adhesive 63) under UV light for 5 min. The coverslips were then treated in a Zepto B (Diener Electronic) plasma oven for 12 min at 30 W under 1 L/h airflow. Immediately after, vesicle mixtures were added to reaction chambers, and after letting the vesicles settle for 4 min, 3 mM CaCl_2_ was added to enhance vesicle fusion on the activated surface. Chambers were incubated for 1 h at 37°C, washed with PBS through serial washes by vigorous pipetting and incubated with 0.1% fatty acid free BSA (Sigma-Aldrich) in protein storage buffer for 30 min. EcadECD was added to a 2 μg/ml final concentration to these chambers and incubated for 45 min at room temperature before changing to pre-warmed imaging medium with serial washes.

### Zebrafish lines and handling

Zebrafish (*Danio rerio*) were handled under a 14 h light/10 h dark cycle^77^. Embryos were raised at 28.5– 31°C in E3 medium and staged as previously described^78^. The following lines were used: WT ABxTL, *Tg(cdh1-tdTomato)xt18*^31^, *Tg(cdh1-mlanYFP)xt17*^31^ and *Tg(actb2:Myl12.1-eGFP;actb2:Utrophin-mCherry)* generated by crossing the preexisting lines *Tg(actb2:Myl12.1-eGFP)* and *Tg(actb2:Utrophin-mCherry)*^21, 34^. Fish were bred in the aquatics facility of IST Austria according to local regulations, and all procedures were approved by the Ethics Committee of IST Austria regulating animal care and usage.

### Cloning of expression constructs

PCR products from plasmids GFP-AHPH-WT (a gift from Michael Glotzer, Addgene plasmid # 68026) and GFP-AHPH-DM (a gift from Alpha Yap, Addgene plasmid # 71368) were subcloned using following primers to create Gateway attB PCR products: GFP-AHPH-WT (5’-GCAGGATCCCATCGATTATGGTGAGCAAGGGCGAG-3’ and 5’-CGTAATACGACTCACTATAGTTTCAAGGCTTTCCAATAGGTTTGTAGCAA-3’), GFP-AHPH-DM (5’-AATACAAGCTACTTGTTCTTTTTGCAGGATCCCATCGATTATGGTGAGCAAGGGCGAG-3’ and 5’-TCTGGATCTACGTAATACGACTCACTATAGTTCTAGAGGCTCAAGGCTTTCCAATAGGTTTGTAGC-3’). cDNA sequence coding for Ftractin (IP3KA_RAT, residues M10 to G52) was codon-optimized and ordered as a gBlocks Gene Fragment (IDT) with attB arms. All products were recombined with pDONR (P1-P2)(Lawson#208) to create entry clones, and further recombined with p3E mNeonGreen (Allelebiotech), p3E mKO2^79^ or p3E polyA (Chien#302), and pCS2-Dest (Lawson #444) to create expression plasmids.

### Embryo microinjections

All embryos were microinjected with 100 pg *lefty1* mRNA at 1-cell stage to induce ectoderm fate. To visualize F-actin and RhoA activities, or to modulate contractility, following mRNAs were additionally injected into 1-cell stage embryos: 60pg *Ftractin-mNeonGreen*, 60 pg of *Ftractin-mKO2*, 80 pg of *GFP-AHPH-WT*^39^, 80 pg of *GFP-AHPH-DM*^40^, 70 pg *constitutively active Myosin Phosphatase* (*CA-Mypt)*^50^ and 3 pg *constitutively active RhoA* (*CA-RhoA)*^41^. To decrease endogenous Ecad amounts, 4 ng *cdh1* morpholino (5’-TAAATCGCAGCTCTTCCTTCCAACG-3’, GeneTools)^30^ was injected at 1-cell stage. Additionally, to visualize F-actin, 0.125 ng Actin protein from rabbit skeletal muscle labeled with TRITC (Cytoskeletal, Inc.) was injected to 1-cell stage embryos. Synthetic mRNAs were produced using the SP6 mMessage mMachine kit (Ambion) and Actin protein was handled according to the manufacturer’s protocol.

### Preparation of embryo cell cultures

30 min before live imaging, embryos were transferred to pre-warmed (28.5-31°C) 0.9x DMEM/F12 medium^23^ (Sigma-Aldrich) supplemented with GlutaMAX (Gibco) and Penicillin-Streptavidin. The blastoderm caps were dissected from the yolk cells at sphere stage with forceps and transferred to 1.5 ml eppendorf tubes using glass pipettes. In the case of inhibitor use, media in the eppendorfs was exchanged to inhibitor-containing media 10 min before cell seeding. All explants were dissociated by gentle tapping and seeded on bilayers covered with control or inhibitor-supplemented media at 29°C.

### Inhibitor treatments

The following inhibitor concentrations in DMEM/F12 media were used: 10 μM for para-nitroblebbistatin (10 mM stock dissolved in DMSO) (Optopharma Ltd.) and 20 nM for 1-Oleoyl lysophosphatidic acid sodium salt (LPA) (5 mM stock dissolved in water) (Tocris). As controls, DMEM/F12 media with or without DMSO (0.1%) were used depending on the solvent of the pharmacological inhibitors.

### Cell confinement by polydimethylsiloxane (PDMS) confiners

In the case of bilayers without EcadECD (except for adherence assay, see below), cells were put under slight PDMS confinement to increase the imaged contact area. Cells were seeded onto bilayers formed on coverslips glued to the bottom of plastic dishes containing a 17 mm round hole, on which a chamber was created by gluing a ring cut from a 15 ml falcon tube. 1:10 PDMS mixtures (Sylgard 184, Ellsworth Adhesives) were prepared as previously described^80^, degassed for 2 min at 2,000 rpm (mix) and for 2 min at 2,200 rpm (defoam) in a mixer/defoamer (ARE-250, Thinky). PDMS was poured onto a wafer and 10 mm round coverslips that were activated by plasma cleaning were pressed onto this mix. The wafer was baked at 95 °C for 15 min and the 16 μm high micropillar-coated coverslips were gently removed from the wafer to be used as confiners. Before use, a confiner was incubated for 5 min with FBS, washed with PBS and kept in culture medium. For imaging, the confiner was placed on a soft pillar attached to a magnetic glass lid, closed on the cells, and kept in place using a magnetic ring underneath the dish during imaging.

### Microscopy

Imaging was performed using microscopes with heating chambers at 29°C. For imaging contact formation, acquisition was started as soon as cells were seeded. For imaging steady contacts, cells were imaged 10-30 min post seeding. Cultures were imaged for ∼ 1 h maximum, and dividing or apoptotic cells were excluded from the subsequent analysis. Most contacts were imaged using an LSM800 equipped with an Airyscan detector and a Plan-APOCHROMAT 63x/1.4 oil objective (Zeiss). For time lapse imaging of Ecad and RhoA biosensors, which showed weaker signal and/or higher photobleaching, Andor Dragonfly 505 equipped with 1x Andor Zyla sCMOS detector and a CFI Apochromat TIRF 60x/NA 1.49/WD 0.13 mm oil objective (Nikon) was used when quantifying intensity changes over time. For all markers, imaging parameters that minimized photobleaching were used for time-lapse analysis (Supp. Fig. 2e). For imaging bilayers, single molecules and FRAP experiments, A TILL Photonics iMic TIRF System equipped with Andor TuCam detection and a 100x/1.49 (Olympus) oil objective was used.

### Data Visualization and Analysis

All micrographs for figures were adjusted for contrast using Fiji^81^. Data for the rest of the analysis were processed raw. Data were plotted using Prism 6 (Graphpad). For sketches and final formatting of figures, Inkscape (Inkscape Project, 2020) was used.

In Supp. Videos 2 and 3, frame averages were made over 2 subsequent frames for better visualization. Temporal visualization in Supp. Video 2 was performed with the Temporal-Color Code plugin in Fiji. Temporal trajectory construction in Supp. Video 3 was performed with the TrackMate plugin^82^ in Fiji, based on a Gaussian fit with an estimated diameter of 0.5 µm. Linking distance of maximum 0.3 µm and gaps of maximum 3 frames were allowed to account for failures to detect Ecad clusters. Randomly-picked trajectories longer than 1 min were shown in the movie.

### FRAP experiments and analysis

To measure the diffusion constant of EcadECD-Cy5 on different bilayer compositions, photobleaching experiments were performed with a frame rate of 2 s per frame. 5 pre-bleach frames were acquired, followed by photobleaching of an area of about 10 µm × 10 µm. Recovery of the signal was analyzed using the frap_analysis program^83^ implemented in MATLAB (The MathWorks, Natick, MA).

FRAP experiments for cellular Ecad were performed using cells obtained from *Tg(cdh1:mlanYFP)* embryos, with a frame rate of 0.5 s per frame. 5 pre-bleach frames were acquired, followed by photobleaching of an area of about 5 µm × 5 µm at the cell contact. A photobleach correction due to the imaging process was performed using an unbleached area of the contact, and the photobleach curve was normalized to the first pre-bleach data point. To obtain the recovery times and immobile fractions, monoexponential functions were fitted to the recovery curves^84^.

### Analysis of Ecad, GTP-RhoA and F-actin average intensities and Myosin-2 mini-filament density

Contact intensity was measured using a custom Python script, by taking ratios of background intensity-subtracted total intensity to total area determined by local thresholding. For rim-to-center ratios, the outer 20% of the contact radius was defined as the contact rim and the rest as the contact center. For measurements over time, GFP-AHPH and Ftractin-mNeonGreen intensity values were first normalized to the maximum intensity for removing injection-based variations between samples.

AHPH expression was detected both diffusely throughout the contact and as cortical foci. These foci were homogeneously distributed at the contact-free interface (Supp. Fig. 2), and once the contact area stabilized, preferentially localized to the contact rim (Fig.1b). Given that such foci were not found using other RhoA biosensors^85^, they were excluded from the average intensity analysis.

To detect Myosin-2 mini-filaments ILASTIK was used^86^. Percentages of total area positive for Myosin-2 signal in the segmented images were determined using a custom Python script. Contact diameters were also estimated from the segmented contact areas, assuming that contacts were generally symmetrical.

### Quantification of adherence

To check for the specificity of cell-bilayer adhesions, either bilayers were prepared without EcadECD or Ecad was knocked down in cells using *cdh1* morpholino. Cells were considered non-adherent when the contact size was below 177 μm^2^ area (∼ 15µm diameter).

### Analysis of colocalization and rim peak distances

Dual-color images were acquired with the confocal mode of LSM800 and used for colocalization analysis. Images were analyzed using Coloc2 plugin in Fiji. Manders’ coefficients M1 and M2 (0 to 1), which give a fraction of overlap between positive pixels, and Pearson’s correlation coefficient (-1 to 1), which gives a value based on the correlation of intensities at two channels, were calculated^87^.

In order to visualize the colocalization, intensity profiles over a 0.3 µm-thick line perpendicular to the rim were plotted together for both channels. Distances to the F-actin rim peaks from GTP-RhoA, Myosin-2 and Ecad peaks were calculated from such profiles.

### Plotting radial intensity profiles

Radial averages of intensity profiles in roughly symmetrical contacts were generated using the transform function in Fiji to rotate an image around its center of mass. The resulting rotations were averaged and radial intensity was plotted by making a line profile along the contact diameter. For plotting radial profiles from multiple cells, profiles were first normalized to contact length, and then to average intensity.

### Analysis of F-actin and Ecad flows

Time-lapse images of Ftractin-mNeonGreen- or Ecad-mlanYFP-expressing cells were used for flow analysis. Unless specified, mature (>10 min post contact initiation) contacts were used for analysis to rule out contact expansion effects and retracting protrusions at the contact rim were excluded from analysis. The built-in Fiji function Multi Kymograph was used to get kymographs along each cell’s diameter. The motion of fluorescent particles within those kymographs was detected using KymoButler, a deep learning automated kymograph analysis software^88^ in Mathematica 12.1 (Wolfram Research, Inc.). BiKymoButler function was used to detect bidirectional tracks with a particle size of 0.3 µm and a minimum duration of 10 s. From these tracks, location and net velocity of particle movements with respect to the center of mass and track durations were calculated. For radial velocities along the contact, multiple kymographs were made with 10° rotations around the center of mass, and net velocities were plotted against their distance from the center of mass. For histograms, 10 tracks from each analyzed cell were pooled together. For analysis of flows around LPA-induced ectopic foci, kymographs of ∼6 µm in length, were constructed around the foci. The net velocity of particle movements were calculated with respect to foci position.

### Analysis of F-actin network density at the contact center

Ftractin-mNeonGreen-labeled F-actin networks, excluding the contact rim, were extracted using SOAX, a software for quantification of biopolymers networks^89^. For time-lapse images, parameters were adjusted for each movie based on inspection of some frames and the corresponding extracted networks; the saved parameters were later on used to batch process the movies. Using a custom Python script, total network lengths were measured with Skan library functions and divided by contact areas to get network density values^90^. Radial network density profile was calculated with the 1 pixel-thick network map as described above.

### Single particle tracking and analysis of Actin lifetime

TRITC-Actin-injected cells were imaged on the Imic TIRF microscope with ∼100 nm pixel size, using a 561-nm laser line with 100 ms exposure. Acquisition intervals of 1 s, 2 s and 3 s were used to capture time lapses of at least 200 frames. Particle detection and tracking were performed using the TrackMate plugin in Fiji, based on a Gaussian fit with an estimated diameter of 0.3 µm. Linking distance of maximum 0.2 µm and gaps of maximum 2 frames were allowed to account for failures to detect particles. Thresholds were adjusted manually for each experiment and tracks were verified by overlaying with the raw data. Average lifetime of Actin at contacts were calculated using a previously described method^91, 92^. Briefly, effective lifetime was obtained by fitting a monoexponential decay function to the lifetime distribution of trajectories obtained from each cell (Supp. Fig. 5a). This value was corrected for photobleaching by using the varying acquisition intervals to obtain a photobleaching constant, on the basis of which a corrected dissociation rate could be calculated (Supp. Fig. 5b,c).

### Quantification of bleb frequency

Standard deviation projections of time-lapse movies of the contact were made using Fiji to define a stable contact zone. Blebs that extended and retracted back to this stable zone were manually counted for different contractility conditions.

### Theoretical modeling of flow-mediated rim accumulation

For any specie with concentration profile *c*(*x, t*) at position x along the contact (in the center-rim axis) and time t, conservation law dictates that ∂_*t*_ *c* = −∂_*x*_ *cv* + *R*(*c*) where *v*(*x*, *t*) is the local velocity and *R*(*c*) local turnover/reaction rates. This simply indicates that local density changes can occur only from velocity gradients or local reactions. The simplest expression for *R*(*c*) is first-order kinetics 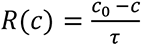, which simply assumes that the specie *c* turns over with a well-defined time scale τ to a target density *c*_0_ (note that both τ and *c*_0_ could in principle depend on time and/or other species, which we neglect here to check whether the simplest model of flow and constant turnover fits the data). The assumption of first-order kinetics was further tested by the fact that it predicts a simple exponential recovery upon FRAP, which fitted well with the data (Fig. 3d’,e). The first of our modeling approaches is to take the velocity field as a given and calculate the resulting profile of specie *c*(*x*, *t*). Based on the data, the simplest expression for velocity is *v*(*x*, *t*) = 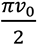 *sin*(2π*x*/*L*), which is zero both at *x* = 0 (center) and *x* = *L*/2 (rim), and has average velocity of *v*_0_ (the experimentally measured parameter, alongside the contact diameter L). This leads - to first order - to a sinusoidal density profile *c*(*x*, *t*) = 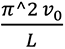 *cos*(2π*x*/*L*), from which we can predict the normalized rim accumulation as 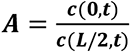 as discussed in the main text. Note that we can also easily incorporate in this model the possibility of immobile fractions, for instance, in Ecad. This can be done by writing two different equations for mobile and immobile Ecad (resp. *c*_*m*_ and *c*_*i*_) ∂_*t*_ *c*_*i*_ = −∂_*x*_*c*_*i*_*v* + *R*(*c*_*i*_). For truly immobile Ecad, we would have *R*(*c*_*i*_) = 0, however this is pathological for long-time scales as all Ecad would rapidly concentrate at the very rim and so as we assume are looking at contacts on the 1-10 min time scale, we assume simply a much second fraction from much slower turnover than the mobile fraction 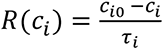, with τ_*i*_ = 3 *min*. This only causes mild increases in rim accumulations of Ecad.

The second step of the modeling is then to ask how the velocity field *v*(*x*, *t*) is determined in the first place. For this, we use the classical isotropic active gel theory, which has been shown to be a minimal description for actomyosin mechanics, and has as input parameters the contractility of actomyosin *χ*, the viscosity of the gel η and the friction to the substrate ξ. The force-balance equation combined with constitutive active gel equation then reads: -ξ *v* = η∂_*xx*_*v* − *χ*∂_*x*_ρ (where ρ is the local actomyosin concentration, also following the conservation equation described above, which has to be complemented with a small diffusion coefficient for stability 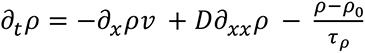). This minimal model has been shown to give rise to spontaneous instabilities, where local accumulations of actomyosin create flows that sustain them despite turnover. Theoretically, this occurs when contractility is above a threshold 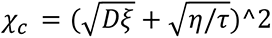. Below the contractility threshold, although spontaneous accumulations are not formed, introducing spatial heterogeneities in the model can lead to strongly self-reinforcing flows. For instance, if we assume that the rim is characterized by slightly larger actomyosin contractility *χ*(1 + δ*χ*) than the center of the contact, this will create flows towards the rim, creating density gradients in ρ which will self-reinforce flows. Interestingly, applied to our data, this suggests that the WT contacts might not be above this threshold (as the patterning is not spontaneous, but instead always strongly guided from the start towards the contact rim) (Supp. Fig. 3e), while the LPA contacts characterized by hyper-activation contractility show the hallmarks of such local and self-organized accumulations (Fig. 2e,e’). Based on our findings that RhoA remains more active at the rim of the contact compared to the center (Fig. 2a), even in the first minutes of contact formation (i.e. prior to flow establishment) (Supp. Fig. 3b), we input in our simulation a higher contractility at the rim, which we take δ=33% under the simple assumption that contractility is proportional to local RhoA activity. Note that in this model, the observed flux of F-actin filaments comprises both diffusion and advection, so we plot in the model the total velocity *v*_*tot*_ = *v* − 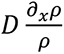. We take the turnover time τ_ρ_ and contact diameter as previously measured. The viscosity η can be used to rescale force balance so we set it to 1 without loss of generality, while the ratio of friction to viscosity ξ/η is taken as ξ/η = 1/100 μ*m*^-2^ based on measurements in other systems^46^, and in order to have flows propagate on the length scale of the entire contact as observed experimentally (Supp. Fig. 3e). The diffusion coefficient - which does not play a key role apart from smoothing the velocity field is taken to be small *D* = 0.1μ*m*^2^/s^64^. Taking a rescaled contractility *χ*/η = 2.25 could reproduce well the average velocity observed in WT contact, as well as the observed velocity and density profile of F-actin as a function of position (Supp. Fig. 6b, also see Fig. 2a and Supp. Fig. 3e). Modeling LPA as moderately larger values for contractility of *χ*/η = 2.75 would result in increased rim to center accumulation, but also in a doubled velocity due to the self-reinforcing loop operating via actomyosin flows, where initial biases in contractility create flows which reinforce the contractility differences between rim and center (Supp. Fig. 6b). Interestingly, in experiments, flows in LPA are comparable in amplitude to the ones measured in WT despite the increased contractility, which can occur if either the friction or viscosity of the actomyosin gel is increased by LPA treatment (e.g. respectively due to stronger link to the substrate or higher cross-linking from Myosin of the active gel - see Supp. Fig. 6b for exploration of the effects of friction or viscosity). Although this value is still below the threshold for purely self-generated instabilities in the absence of any guiding cues, considering local and stochastic increases in contractility can lead to local flows and actomyosin accumulation^64^, reminiscent of the local foci that we sometimes observe under LPA treatment (Fig. 2e). Interestingly, localized accumulations have been recently predicted in similar models of active cytoskeleton additionally considering the dynamics of activators^93^, which would be an interesting avenue of investigation for the future.

### Statistics and Reproducibility

Statistical tests were performed in Prism 6. Details for each experiment are described in figure legends. In brief, a D’Agostino-Pearson normality test was first performed, and, based on the results, a two-tailed Student’s t-test for parametric distributions and a Mann-Whitney *U*-test for non-parametric distributions were used to compare two groups. To compare more than two groups, an ANOVA test for parametric distributions and a Kruskal-Wallis test for non-parametric distributions were used. Independent experiment N, when involving cells, denotes a single embryo, where controls and experiments were performed on the same day and within the same egg-lay. n denotes the number of cells analyzed.

## Data and Code Availability

All data and code supporting the current study are available from the corresponding author upon request.

## Acknowledgements

We thank members of the Heisenberg and Loose Labs for their help and feedback on the manuscript, and the Aquatics and Imaging & Optics facilities of ISTA for their continuous support. This work was supported by an ERC Advanced Grant (MECSPEC) to C.-P.H.

## Author Contributions

F.N.A. and C.-P.H. designed the research. F.N.A. performed the experiments and analyzed the experimental data. E.H. performed numerical simulations and modeling. J.M. designed and produced the wafers for PDMS confiners. M.L. provided reagents, conceptual input and support with analysis. F.N.A and C.-P.H. wrote the manuscript. All authors edited the manuscript.

## Declaration of Interests

The authors declare no competing interests.

## Notes

### Competing Interest Statement

The authors have declared no competing interest.

